# Knowledge-driven perceptual organization reshapes information sampling via eye movements

**DOI:** 10.1101/2021.09.24.461220

**Authors:** Marek A. Pedziwiatr, Elisabeth von dem Hagen, Christoph Teufel

## Abstract

Humans constantly move their eyes to explore the environment and obtain information. Competing theories of gaze guidance consider the factors driving eye movements within a dichotomy between low-level visual features and high-level object representations. However, recent developments in object perception indicate a complex and intricate relationship between features and objects. Specifically, image-independent object-knowledge can generate objecthood by dynamically reconfiguring how feature space is carved up by the visual system. Here, we adopt this emerging perspective of object perception, moving away from the simplifying dichotomy between features and objects in explanations of gaze guidance. We recorded eye movements in response to stimuli that appear as meaningless patches on initial viewing but are experienced as coherent objects once relevant object-knowledge has been acquired. We demonstrate that gaze guidance differs substantially depending on whether observers experienced the same stimuli as meaningless patches or organized them into object representations. In particular, fixations on identical images became object-centred, less dispersed, and more consistent across observers once exposed to relevant prior object-knowledge. Observers’ gaze behaviour also indicated a shift from exploratory information-sampling to a strategy of extracting information mainly from selected, object-related image areas. These effects were evident from the first fixations on the image. Importantly, however, eye-movements were not fully determined by object representations but were best explained by a simple model that integrates image-computable features and high-level, knowledge-dependent object representations. Overall, the results show how information sampling via eye-movements in humans is guided by a dynamic interaction between image-computable features and knowledge-driven perceptual organization.

## Introduction

Human visual experience carves up the world into objects (Feldman, 2003; Wagemans et al., 2012), distinct entities that are critical in structuring our interaction with the environment. When searching for a specific item in a scene or when exploring the world with no purpose other than to obtain information, humans tend to look at the centre of objects (e.g., Nuthmann & Henderson, 2010; Pajak & Nuthmann, 2013; Stoll, Thrun, Nuthmann, & Einhäuser, 2015). While these object-oriented effects of information sampling are well established, the current literature provides little consensus about which specific aspects of objects influence programming of eye movements (Borji & Tanner, 2016; Federico & Brandimonte, 2019; Henderson, Malcolm, & Schandl, 2009; Kilpelaïnen & Georgeson, 2018; Nuthmann, Schütz, & Einhäuser, 2020; Van der Linden, Mathôt, & Vitu, 2015). Current explanations of oculomotor control either highlight the importance of image-computable features that are correlated with object locations (Adeli, Vitu, & Zelinsky, 2017; Borji & Tanner, 2016; Russell, Mihalaş, von der Heydt, Niebur, & Etienne-Cummings, 2014) or emphasize the role of high-level objecthood and semantics (Hayes & Henderson, 2021; Henderson & Hayes, 2017; Henderson et al., 2009; Hwang, Wang, & Pomplun, 2011; Nuthmann & Henderson, 2010; Stoll et al., 2015; Melissa Le Hoa Võ, 2021). However, disentangling the different aspects of objects and identifying their importance for eye-movement control is complicated by the intricate and complex relationship between image-computable features and object representations: features are necessary for visual object representations to arise – hallucinations being an exception (Horga & Abi-Dargham, 2019; Powers, Mathys, & Corlett, 2017; Teufel et al., 2015) – but they are often not sufficient. Indeed, a growing number of studies using human psychophysics (Christensen, Bex, & Fiser, 2015; Lengyel, Nagy, & Fiser, 2021; Lengyel et al., 2019; Neri, 2017; Ongchoco & Scholl, 2019; Teufel, Dakin, & Fletcher, 2018) neuroimaging (Flounders, González-García, Hardstone, & He, 2019; Hsieh, Vul, & Kanwisher, 2010), and animal electrophysiology (Gilbert & Li, 2013; Liang et al., 2017; Self et al., 2019; Self, van Kerkoerle, Supèr, & Roelfsema, 2013; Walsh, McGovern, Clark, & O’Connell, 2020) suggest that in order for object representations to emerge, prior object-knowledge has to interact with sensory processing. By contrast to conventional models of object recognition (DiCarlo, Zoccolan, & Rust, 2012; Kourtzi & Connor, 2011; Kriegeskorte, 2015; Marr & Nishihara, 1978), these studies demonstrate that prior object-knowledge effectively generates objecthood by reconfiguring sensory mechanisms that process visual inputs, thereby changing how feature space is carved up into meaningful units (Teufel & Fletcher, 2020). In the current study, we abandon the simplifying dichotomy between image-computable features and high-level object representations. We show that the dynamic re-shaping of feature space by knowledge-driven perceptual organization has a substantial influence on information sampling via eye movements in human observers.

The most influential early models of eye-movement control largely disregarded objects, arguing that programming of eye-movements is determined by an analysis of low-level features such as luminance, colour, and orientation (Harel, Koch, & Perona, 2007; Itti & Koch, 2001). According to these early accounts, the visual system computes feature maps, which highlight areas in the image that attract fixations (Zelinsky & Bisley, 2015). Over the past 15 years, however, several studies have emphasised the importance of objects and semantic meaning in guiding information sampling (Einhäuser, Spain, & Perona, 2008; Henderson & Hayes, 2017; Hwang et al., 2011; Nuthmann & Henderson, 2010; Pajak & Nuthmann, 2013; Pilarczyk & Kuniecki, 2014; Rider, Coutrot, Pellicano, Dakin, & Mareschal, 2018; Stoll et al., 2015). For instance, in one of the early studies, Einhäuser and colleagues (2008) found that maps of object locations outperform maps derived from a low-level feature model in predicting human fixations. Moreover, human observers show a tendency to look at the centre of objects rather than their edges, contrasting with predictions from early low-level feature models (Nuthmann & Henderson, 2010; Pajak & Nuthmann, 2013; Stoll et al., 2015; see also Vincent, Baddeley, Correani, Troscianko, & Leonards, 2009). These effects have been interpreted as demonstrations of the importance of objects in oculomotor control. An even more ambitious approach is based on a novel technique called meaning maps (Henderson & Hayes, 2017). Such maps are created by segmenting a visual scene into small, isolated patches, which are rated for their meaningfulness. These ratings are pooled together into a smooth map, which is assumed to capture the distribution of meaning across an image. A key paper adopting this approach showed that meaning maps are better at predicting human fixations in comparison to GBVS (Harel et al., 2007), a low-level feature model. These results have been interpreted as evidence to suggest that eye-movements are controlled by the semantic properties of images (Henderson & Hayes, 2017).

The notion that eye-movements are controlled by high-level object representations, or by the meaning of image parts has not remained (Borji et al., 2013; Elazary & Itti, 2008; Kilpelaïnen & Georgeson, 2018; Masciocchi et al., 2009; Wu et al., 2014). For instance, a recent attempt to assess the unique contribution of features vs. objects to oculomotor control suggests that object-centred effects are, at least partly, driven by low-level features that correlate with objects (Nuthmann et al., 2020). This conclusion is in line with a careful psychophysical study, suggesting that the tendency of human observers to focus on the centre of objects might be controlled by a relatively simple process that programs eye-movements towards homogeneous luminance surfaces on the basis of luminance-defined edges (Kilpelaïnen & Georgeson, 2018). This result provides a potential mechanism for the finding that fixations that occur shortly after image onset tend to be located close to the stimulus centre not only for objects but also for non-objects if low-level properties are matched (Van der Linden et al., 2015). Together, these results suggest that the tendency to fixate on the centre of objects might not be related to objecthood itself but is controlled by mechanisms that respond to relatively low-level features in the input. Note, however, that the study by van der Linden and colleagues (2015) also suggests that guidance of eye-movements that are generated later after image onset might be affected by semantic aspects of object. This finding potentially indicates a time course according to which locations of early fixations are mainly determined by low-level, image-computable features while locations of later fixations might be determined by genuine object representations (see also Anderson, Ort, Kruijne, Meeter, & Donk, 2015).

More generally, a limitation of many previous studies that aim to show the contribution of objects, or semantic meaning to oculomotor control is their reliance on a comparison to computational models that calculate image-computable feature maps as their null hypothesis. The typical approach (Pedziwiatr, Kümmerer, Wallis, Bethge, & Teufel, 2021b) of these studies is (i) to compute a saliency map based on certain features of images used in the experiment, (ii) to generate a map of semantically important regions or object locations in these images, and (iii) to assess which of the two maps better predicts human fixations (for example, see Henderson & Hayes, 2017; Pilarczyk & Kuniecki, 2014; Rider et al., 2018). Insofar as one of these two maps better predicts human fixations, that factor is considered to be critical in gaze guidance. This approach has led to important insights regarding oculomotor control but is hampered by its dependence on saliency models, with the specific choice of model being critical in determining the interpretation (Borji et al., 2013; Pedziwiatr et al., 2021b; Pedziwiatr, Kümmerer, Wallis, Bethge, & Teufel, 2021a). For instance, the conclusion that objects *per se*, rather than image-computable features that are correlated with objects, guide human eye-movements, as suggested by Einhäuser and colleagues (2008), was based on a comparison between manually labelled objects maps and a map derived from one of the earliest saliency models (Itti & Koch, 2000). A re-analysis of the data compared the object maps to other low-level models, including the AWS model (A. Garcia-Diaz, Leboran, Fdez-Vidal, & Pardo, 2012; Antón Garcia-Diaz, Fdez-Vidal, Pardo, & Dosil, 2012), which outperformed the high-level object model in predicting human performance (Borji et al., 2013). This finding thus led to a reversal of the original conclusion. When a very similar dataset was analysed for a third time (Stoll et al., 2015), a more sophisticated object map showed higher performance than even AWS. Note, however, that since publication of these studies, saliency models that outperform AWS by a large margin have been developed (Kümmerer, Wallis, Gatys, & Bethge, 2017; Thomas, 2016), and yet another re-analysis might show that these novel models outperform even the more sophisticated object maps. This string of studies illustrates the idea that the conclusions are determined by the choice of model. A similar dispute regarding the usefulness of meaning maps serves as another example: the seminal meaning maps study claimed that eye-movements are driven by meaning rather than image-computable features because meaning maps were better at predicting fixations than one specific saliency model – the GBVS model (Harel et al., 2007). However, subsequent work showed that meaning maps are outperformed by more advanced saliency models that are based on image-computable features, such as DeepGaze II (Kümmerer, Wallis, Gatys, Bethge, et al., 2017), challenging the initial interpretation (Henderson, Hayes, Peacock, & Rehrig, 2021; Pedziwiatr et al., 2021a, 2021b).

Independently of the favoured interpretation of these findings, there is a more fundamental aspect that is easily overlooked. The emphasis on a dichotomous view, which contrasts outputs of low-level feature models with ‘objects’ or ‘semantic information’, and the tendency to conceptualise these as categorically different interpretations, has concealed a fundamental similarity between these explanations. Specifically, comparable to how low-level models deal with simple features, most studies implicitly treat ‘objects’ or ‘semantic information’ as image-computable properties. This notion is also the basis for state-of-the-art computer vision models that aim to predict human fixations (e.g., Kroner, Senden, Driessens, & Goebel, 2020; Kümmerer et al., 2017a): these models use deep convolutional neural networks trained on object recognition to extract high-level features that are directly computed from the image. In other words, rather than providing diametrically opposed interpretations, the different approaches in the current eye-movement literature can be understood as lying on a continuum, with their position being defined by the features they emphasise. This notion is made explicit in a recent study by Schütt and colleagues (Schütt, Rothkegel, Trukenbrod, Engbert, & Wichmann, 2019): the authors explicitly conceptualised objects as high-level features that are computed in a bottom-up fashion, and contrasted their contribution to the guidance of eye-movements with the contribution of low-level features.

While the theoretical precision of the study by Schütt and colleagues is exceedingly helpful in clarifying the different positions, conceptualising objects as high-level features directly conflicts with current developments in object perception. Two aspects of the complex relationship between features and objects are particularly relevant: first, several recent studies demonstrate that features are not always sufficient for object representations to arise (Flounders et al., 2019; Hsieh et al., 2010; Lengyel et al., 2019, 2021; Ongchoco & Scholl, 2019; Teufel et al., 2018). Rather, objecthood emerges as a consequence of the interaction between current visual input and perceptual organization processes that are based on prior object-knowledge. Second, once object representations have been generated, top-down influences reconfigure the way in which even some of the earliest cortical mechanisms process low-level visual features (Christensen et al., 2015; Flounders et al., 2019; Hsieh et al., 2010; Lengyel et al., 2021, 2019; Neri, 2014, 2017; Ongchoco & Scholl, 2019; Teufel et al., 2018). For instance, psychophysical studies show that early feature-detector units are sharpened for currently relevant input based on top-down influences from object representations (Teufel et al., 2018). This reconfiguration of information processing is detectable in early retinotopic cortices (Flounders et al., 2019; Hsieh et al., 2010). Overall, these findings thus cast serious doubt on the notion that the human visual system computes image features independently of the inferred object structure of the environment (Neri, 2017).

This novel perspective of object perception has fundamental implications for our understanding of information sampling via eye movement. First, if objecthood emerges from the interaction between features and prior knowledge, then the question of whether objects guide eye movements cannot be answered by an approach that exclusively focuses on how image-computable feature space is carved up by the visual system, regardless of whether the considered features are low- or high-level. Second, the novel perspective of object perception means that a full understanding of the role of objects in eye-movement control has to move away from regarding feature space as static, instead taking into account the plasticity of low-level sensory processing introduced by dynamic interactions with object representations. Here we address both of these issues. We analysed gaze data from human observers viewing stimuli, which, on initial viewing, are experienced as a collection of meaningless black and white patches. After gaining relevant object knowledge, however, the observers’ visual system organizes the sensory input into meaningful object representations. We demonstrate that this knowledge-driven perceptual organization of identical inputs substantially re-shapes eye-movement patterns, with the selection of fixation locations being driven by a combination of image-computable features and the knowledge-dependent object representations. Moreover, these effects are already present at the first fixation. In summary, we show that a fundamental human visual behaviour – information sampling via eye movements – is guided by a dynamic interaction between image-computable features and object representations that emerge when prior object-knowledge restructures sensory input.

## Experiment 1 – Methods

### Overview

In Experiment 1, observers viewed black and white two-tone images while their eye movements were recorded. Two-tone images are derived from photographs of natural scenes (‘templates’). Each two-tone appears as meaningless patches on initial viewing. Once an observer has acquired relevant prior object-knowledge by viewing the corresponding template, however, processes of perceptual organization in the visual system bind the patches of the two-tone image into a coherent percept of an object (see caption of Fig. 1 for instructions of how to experience the effect).

**Fig. 1.**
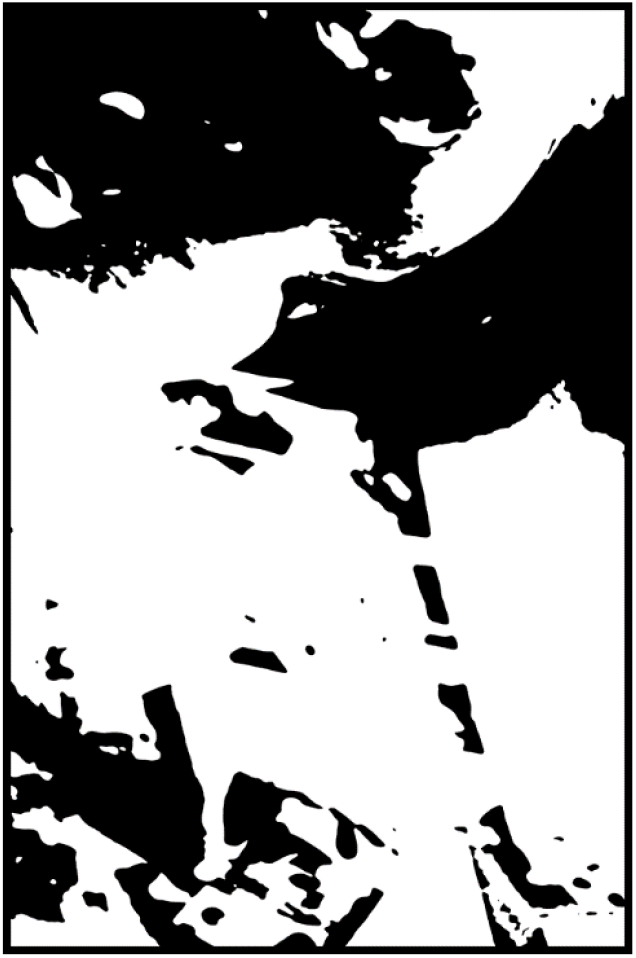
Example of a two-tone image. On initial viewing, this image appears as meaningless black and white patches. To be able to perceptually organize it into a meaningful percept, the reader is advised to first carefully look at the template image from which this two-tone was derived, presented on Fig. 2. Note that the example two-tone image is for illustration only, it was not used in the study. Image copyrights owner: author C. T.

Two-tone images provide a tool to manipulate object perception without changing the visual features of the stimulus. They are therefore ideally suited to test the hypothesis that human oculomotor control is determined by object representations that are not constituted by image-computable features but emerge via an interaction between image-computable features and prior object-knowledge. According to this idea, eye movements in response to two-tone images should be influenced by whether the observer experiences the input as an object percept. Specifically, patterns of fixations on identical two-tone images should be more similar to the ones from the corresponding template when an observer experiences the two-tone image as a meaningful object percept compared to when they experience it as meaningless patches.

To test these predictions, we recorded eye-movements of 36 human observers who viewed two-tone images before and after being exposed to the relevant templates (Before, After, and Template conditions, respectively; see Fig. 2). In the Before condition, observers perceive two-tone images as meaningless black and white patches. In the After condition, prior object-knowledge allows them to bind patches into meaningful object percepts. Crucially, any potential differences in eye movements between the Before and the After conditions cannot be explained by image-computable features because these are identical across these conditions; the only aspect that has changed is the prior object-knowledge that observers have access to. Experiment 1 established the key effects; to exclude alternative explanations, we conducted Experiments 2 and 3 (see Fig. 3 for design details).

**Fig. 2.**
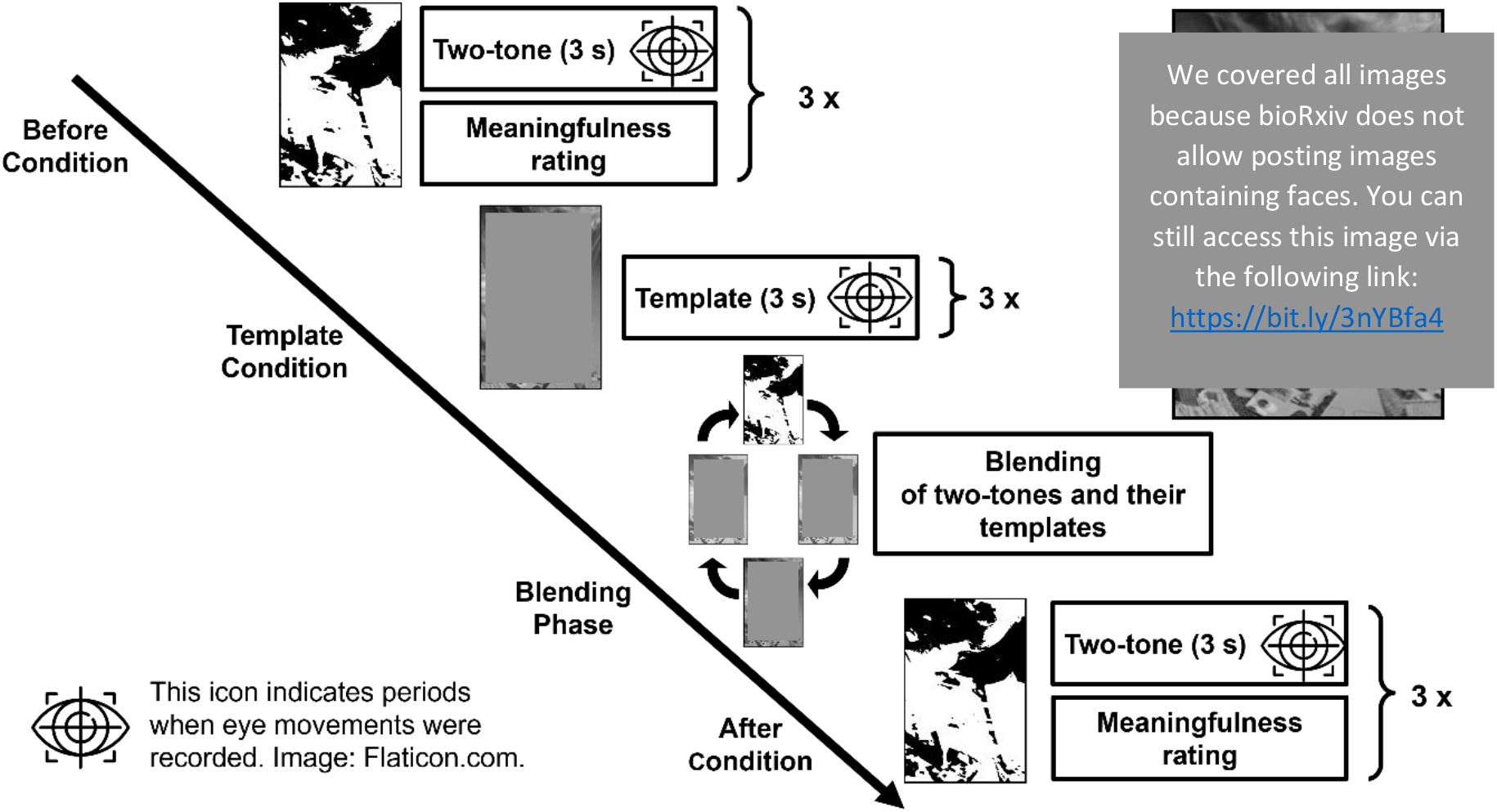
Experiment 1 – Outline of a single experimental block. In each block, observers first free-viewed three two-one images (Before condition). After presentation of each image, they were asked to rate its perceived meaningfulness by adjusting a visual analogue scale (i.e., a ‘slider’). This task was included as a manipulation check. Next, the grayscale templates of these three two-tones were presented (Template condition). In order to ensure that observers acquired the relevant object-knowledge essential to bind the two-tone image into a meaningful percept, in the next part of the block, observers viewed the two-tones gradually blended with their templates six times (Blending Phase). The After condition was identical to the Before condition in all aspects except for the order of presentation of the two-tone images. The whole experiment consisted of 10 blocks (30 two-tones in total) and throughout each block, the eye movements of observers were recorded. In the upper right corner, the template of the two-tone image from Fig. 1 is presented (copyrights owner: author C. T.).

**Fig. 3.**
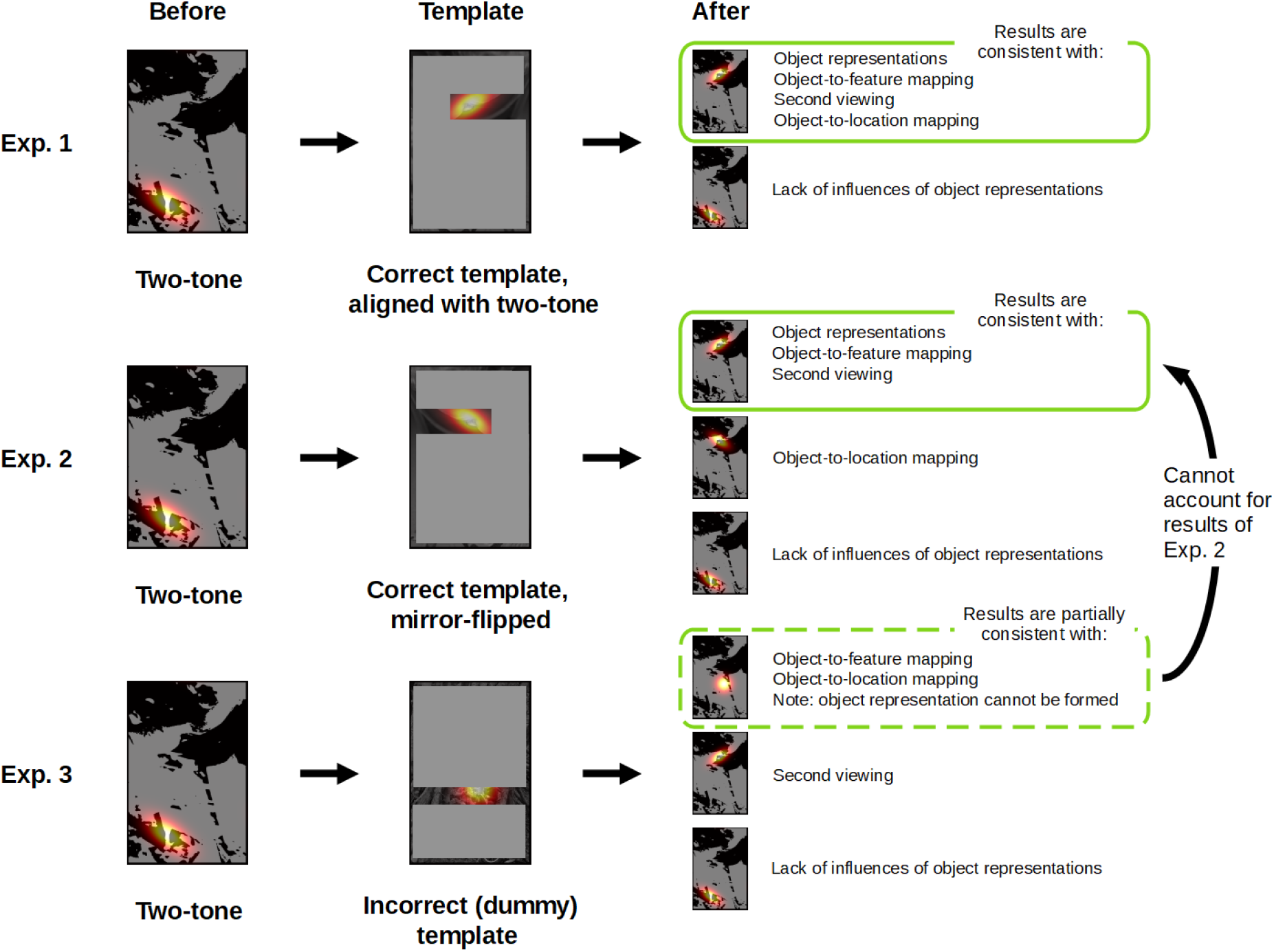
Summary of key experimental manipulations, predictions, and findings of Experiment 1, 2, and 3. The heatmaps superimposed over the example stimuli illustrate different hypotheses about the role of knowledge-driven perceptual organization in gaze guidance. The green rectangles indicate the hypotheses that are consistent with the results of each experiment. All three experiments had a similar design: eye-movements were recorded while observers first viewed a two-tone image without prior knowledge in the Before condition (left column), were then presented with a grayscale image in the Template condition (middle), and finally saw the same two-tone image again in the After condition (right). The key difference across the three experiments was what type of grayscale image was shown in the Template condition. In Experiment 1, the original grayscale photograph used to generate the two-tone image was shown. This provided the observers with the prior object-knowledge required to organize the two-tone image into a coherent object percept in the After condition. If this knowledge-driven perceptual organization does not affect gaze guidance, eye-movements in in the After condition should be identical to those in the Before condition (bottom panel in first row/last column). By contrast to this prediction, we found that gaze guidance in the After condition was similar to that in the Template condition (top panel in first row/last column). This finding is consistent with the idea that knowledge-driven perceptual organization is an important driver of oculomotor control. However, it is also consistent with three alternative interpretations, which we excluded in Experiments 2 and 3. In Experiment 2, the template was the original grayscale photograph, from which the two-tone had been generated, but by contrast to Experiment 1, it was mirror-flipped. This manipulation provided the means to exclude the hypothesis that when viewing the template, observers learned the position of objects in the images, and re-visited these locations in the After condition. In Experiment 3, instead of the correct templates of the two-tones images, ‘dummy templates’ which were unrelated to the two-tone images were presented. This final experiment allowed us to assess the extent to which findings in Experiment 1 and 2 might be explained simply by the fact that in the After condition a two-tone image was viewed for the second time, or that observers had learned to map the features of a two-tone image to locations of objects in the template images. The findings are not consistent with the second-viewing hypothesis. However, we found a small effect consistent with the feature-to-object mapping hypothesis. Importantly, while this might be a contributing factor to the results of Experiment 1 and 2, we demonstrate that this effect is far too small to fully account for the main findings.

### Observers

The primary units of analysis were not individual observers, but the distribution of fixations from all observers on individual images. Therefore, we selected the number of observers based on the estimation of how well our empirical fixation distributions approximate the theoretical distributions which would be obtained from the population of infinitely many observers.

Previous work has shown that fixations from 18 observers provide a sufficiently good approximation for natural scenes, and that further increasing the number of observers results only in marginal improvements (Judd et al., 2012). However, one of our analyses required splitting our sample into two groups and we therefore recruited 36 observers in total, ensuring sufficient amounts of data in each group after the split. All participants were Cardiff University students, had normal or corrected-to-normal vision, participated in the study voluntarily, and received either money or study-credits as a reimbursement. All experiments reported in this article were approved by the Cardiff University School of Psychology Research Ethics Committee.

### Stimuli

We used 30 pairs of images, where each pair consisted of a two-tone image and its template in greyscale. These stimuli were a subset of stimuli used in a previous study (Teufel et al., 2015), where details of template selection and two-tone image generation can be found. In brief, template images – predominantly photographs of animals in their natural environments – were taken from the Corel Photo library. Two-tones were generated by smoothing and binarising template images. A good two-tone image should be perceived as a collection of meaningless patches prior to seeing its template but observers should be able to easily bind the stimulus into a coherent percept of an object after they see the template. Extensive tests on naïve observers were conducted to select both the template images, and the parameters of smoothing and binarisation that guarantee that the created two-tones have these desired properties.

### Experimental setup

The experiment was conducted in a dark testing room. Participants sat 56 cm from the monitor, with their head supported by a chin and forehead rest. Their eye-movements were recorded using an EyeLink 1000+ eye-tracker (with 500 Hz sampling rate) placed on a tower mount. The experiment was controlled by in-house developed code written in Matlab R2016b (Mathworks, Natick, MA) and using the Psychophysics Toolbox Version 3 (Brainard, 1997; Kleiner et al., 2007). Images were presented centrally on the screen, against a mid-grey background. Images measured 21.9o visual angle (788 pixels) horizontally and 14.6o (526 pixels) vertically.

### Procedure

The experiment consisted of ten blocks, a single block is schematically illustrated in Fig. 2. Before the start of the procedure, a 13-point eye-tracker calibration and validation was conducted. Each block started with the Before condition, in which three two-tones were presented in a sequence, each for 3 seconds. Observers were instructed to carefully look at these images. Two-tone images were preceded by a centrally-located fixation-dot displayed for 1 second. They were followed by a visual analogue scale, which observers adjusted by pressing ‘z’ and ‘m’ buttons on a keyboard to indicate how meaningful they experienced the two-tone image to be. Meaningfulness ratings were used as a manipulation check. After each rating, a blank screen was displayed for 500 ms. The Before condition was followed by the Template condition, in which template images were displayed while eye-movements were recorded – again, each for 3 seconds, preceded by a fixation dot. After the Template condition, we ensured that observers had enough object-knowledge to bind two-tone images into meaningful object percepts by presenting six cycles of dynamic blending between two-tones and their templates (Blending Phase). Each cycle began with the presentation of a template image for two seconds. This was then linearly blended into the corresponding two-tone image, with the full transition from template to two-tone taking 4 seconds. The two-tone image remained on the screen for 2 seconds and then was blended back into the template, remaining on the screen for another 2 seconds. Each of the three image-pairs used in a block was presented in a full blending procedure twice with the order pseudorandomised such that the same pair was never used twice in a row. The subsequent cycles of blending were separated with a blank screen presented for 500 ms. After the Blending Phase, the After condition was presented, which was identical to the Before condition except that images were presented in a newly randomized order. There was a break every two blocks, and the eye-tracker was re-calibrated. For each observer, images were assigned to blocks randomly and were presented in a pseudo-random order within each block. The pseudo-randomization ensured that the image shown last in the Blending Phase was never presented at the beginning of the After condition. Total experiment time was ∼50 minutes.

Instructions were delivered verbally and on-screen. Key elements of the procedure were illustrated visually: observers were shown a single two-tone image (which was not used in the actual experiment), rated its meaningfulness, viewed the blending procedure with the template and, finally, viewed the same two-tone again and were asked to provide a meaningfulness rating.

### Data pre-processing and analysis methods

The default EyeLink algorithm was used to extract fixation-locations from the eye-movement recordings. Further data pre-processing was done in Matlab. For each image, we discarded the initial fixation that was directed at the fixation-dot presented before image onset. We also discarded fixations not landing within the image-boundaries. Further details regarding data exclusions can be found in the *Data exclusion* section of the Supplement. For each image in each condition, we generated heatmaps (see examples on Fig. 5E) by smoothing the discrete distribution of fixations with a Gaussian filter, cutoff frequency of –6dB (implementation provided by Bylinskii and colleagues; Kümmerer et al., 2020), and then normalizing the smoothed distribution to the zero-one range.

The majority of our analyses focused on the similarity between two heatmaps. As a similarity index, we calculated Pearson’s linear correlation coefficient using Matlab implementation (Kümmerer et al., 2020). This measure is intuitive, commonly used in the literature (Wilming, Betz, Kietzmann, & König, 2011), and its values have a straightforward interpretation. In the current study, values ranged between zero and one, with one indicating that two heatmaps are identical and zero indicating a maximal dissimilarity. In the Supplement, we provide the results of key analyses using a different metric to quantify the similarity between two heatmaps, the histogram intersection (SIM; Bylinskii, Judd, Oliva, Torralba, & Durand, 2016), showing a similar pattern of results. For statistical comparisons, we primarily relied on standard null-hypothesis-significance-testing techniques implemented in R (R Core Team, 2020) and Matlab. Unless otherwise stated, the t-tests reported throughout the text are paired-sample t-tests. In order to assess the amount of evidence for a lack of a difference between groups of measurements, we used Bayes factors (BFs) calculated using bayesFactor R package (Morey & Rouder, 2018).

## Experiment 1 – Results

### Manipulation check: prior object-knowledge changes perceived meaningfulness of two-tone images

In the Before and After conditions, observers rated the perceived meaningfulness of two-tone images. Averaging these ratings per image showed that the two-tones were perceived as more meaningful in the After compared to the Before condition (Fig. 4A and B; t(29) = 23.84, p < 0.001; mean difference M_diff_ = 0.36, 95% confidence interval CI = [0.33, 0.4]). The same pattern of results held when the ratings were averaged per observer (t(35) = 14.42, p < 0.001; M_diff_ = 0.37, 95% CI = [0.31, 0.42]). These results provide a manipulation check, suggesting that observers are able to organize two-tone images into meaningful object representations after but not before acquiring relevant prior object-knowledge.

**Fig. 4.**
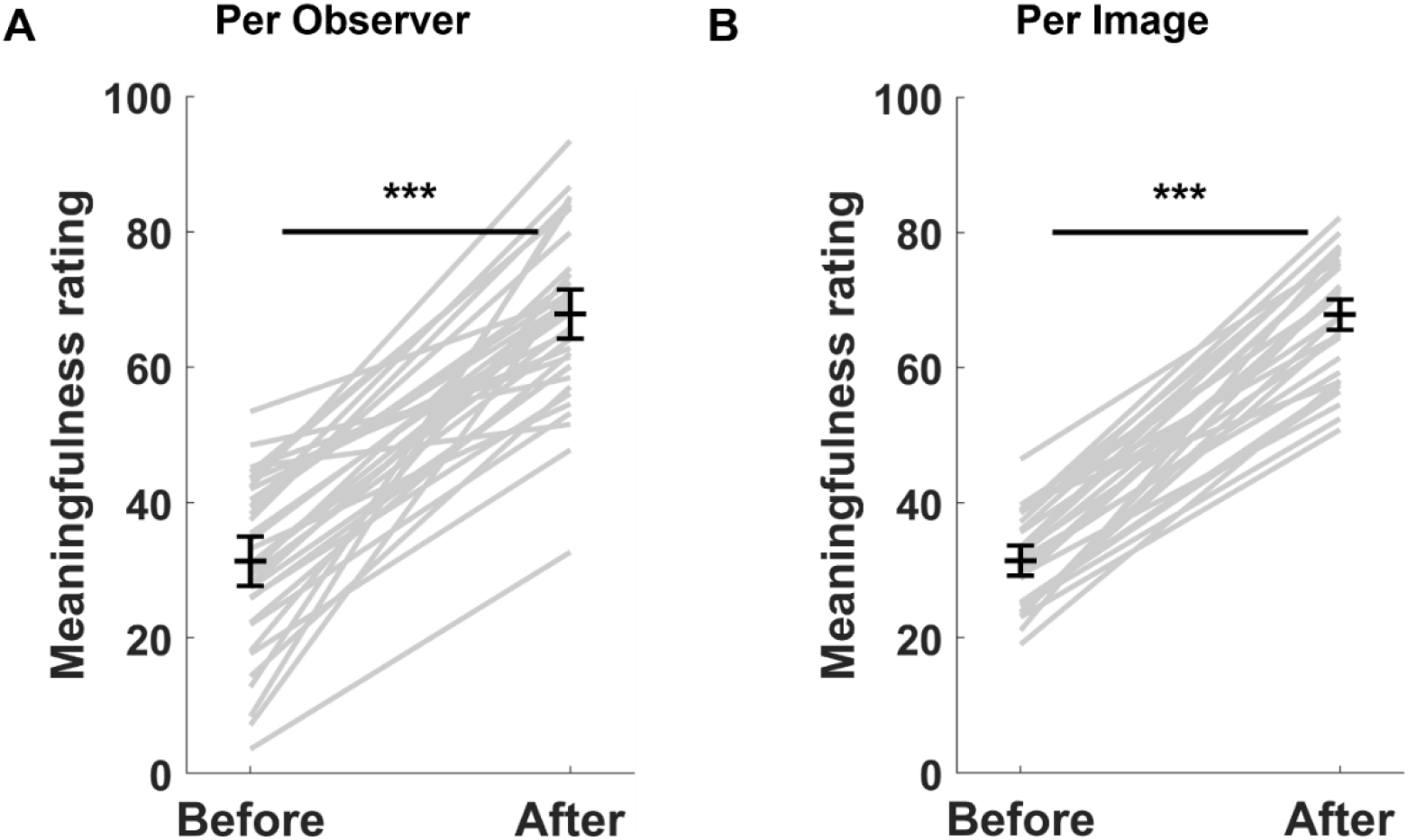
Meaningfulness ratings for two-tone images in the Before and After conditions. As expected, two-tone images were rated as more meaningful in the After than the Before condition, both when the ratings were averaged per observer (A) and per image (B). This finding demonstrates that acquiring relevant object-knowledge changes perception. The following conventions are used in this and all remaining figures: asterisks on plots indicate p-values: *** indicates p ≤ 0.001, ** indicates p ≤ 0.01, * indicates p ≤ 0.05, and ‘n.s.’ indicates the lack of statistical significance. Grey lines indicate values for individual observers (panel A) and images (panel B). Black horizontal bars indicate means. They are surrounded with 95% confidence intervals for within-subjects designs, calculated using Cousineau-Morey method (Cousineau, 2005; Morey, 2008).

### Knowledge-dependent object representations control the spatial distributions of fixations

If knowledge-dependent object representations drive eye movements, the spatial distribution of fixations recorded in response to two-tone and template images should be more similar when two-tone images elicit object representations (After condition) compared to when they do not (Before condition). To test this hypothesis, we compared the similarities of heatmaps across pairs of conditions (Fig. 5A). As predicted, we found a higher similarity between the Template-After pair (M = 0.90, SD = 0.07) compared to the Template-Before pair (M = 0.72, SD = 0.13; t(29) = 8.39, p < 0.001; mean difference M_diff_ = 0.18, 95% CI = [0.14, 0.22]). This result suggests that gaze patterns in response to two-tone images more closely resemble eye movements from the templates when the two-tones were perceived as containing meaningful objects, as compared to when they were perceived as meaningless patches.

**Fig. 5.**
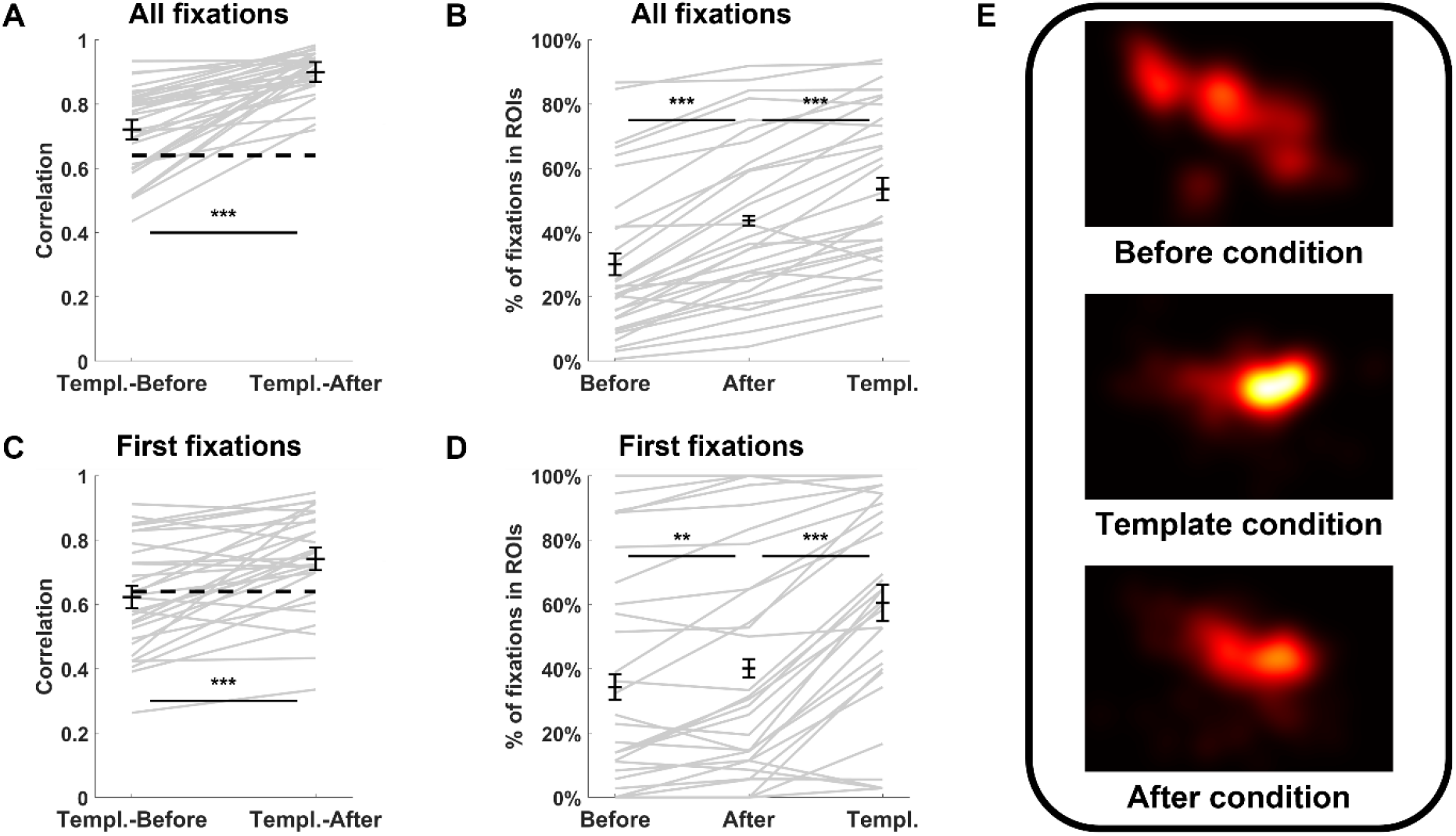
Results of Experiment 1. A) Similarities between heatmaps from template and two-tone images, where the two-tones were viewed either in the Before or in the After condition. When observers organized the two-tones into coherent percepts of objects, their fixation patterns on two-tone images became more similar to the gaze patterns registered on the corresponding template images. Heatmaps illustrating this effect for one image are shown on panel E. The dashed horizontal line illustrates the baseline, i.e., the expected similarity with the Template condition based purely on centre bias. B) Percentage of fixations landing within the regions-of-interest (ROIs) in each condition. ROIs included important object parts (e.g., the heads of depicted animals). On the same two-tone image, ROIs attracted more fixations in the After than in the Before condition, consistently with the hypothesis that in the After condition eye-movements were guided by object representations. C, D) The same analyses as on panels A and B but conducted including only first fixations from the Before and After conditions. In both analyses, the effects of knowledge-driven perceptual organization were still evident. E) Sample heatmaps illustrating the distributions of fixations in all three conditions of Experiment 1 for one two-tone/template pair. These maps were created from all fixations registered on the images. Pixel values of all three maps were jointly normalised to zero-one range, so colour values (indicating fixation densities) are comparable across panels.

While there was a clear difference in similarity between the two pairs, at first glance the Template-Before similarity might seem unexpectedly high. Importantly, however, the distribution of fixations on images is not only determined by the characteristics of the visual input but also by general factors that are independent of the image (Tatler & Vincent, 2009). One key general factor is the centre bias, a tendency of humans to look at the centre of an image rather than regions closer to the edges (Tatler, 2007). A meaningful evaluation of the difference in similarities between Template-Before and Template-After pairs therefore requires a baseline that accounts for this bias. We modelled a centre bias for our data by creating a single heatmap (labelled ‘Centre’) from all fixations registered throughout the experiment. We found a statistically robust difference in similarity scores between the Template-Centre and Template-Before pairs (Template-Centre: M = 0.64, SD = 0.16; Template-Before: M = 0.72, SD = 0.13; t(29) = 2.40, p = 0.023; M_diff_ = 0.08, 95% CI = [0.01, 0.14]). Importantly, however, this difference was small, suggesting that a centre bias explained most, but not all, of the Template-Before similarity.

We ran a further analysis (full details in Supplement) to address the influence of knowledge-dependent object representations by comparing heatmaps from identical visual inputs only. In other words, instead of analysing the similarity between heatmaps from a two-tone image and its template image (different visual inputs), we evaluated the similarities in heatmaps when the same two-tone image was viewed in the Before and the After conditions (identical visual inputs). The findings provide further support for the influence of object-knowledge on gaze guidance (see supplement for details).

### ROI analysis: changes to the spatial distribution of fixations are specific to object-related image content

The analyses of heatmap similarities suggests that prior object-knowledge contributes to eye-movement control. We used a region-of-interest (ROI) analysis to assess in a more fine-grained manner the extent to which changes in fixation patterns related directly to object representations. We exploited the fact that animal and human heads are known to attract fixations in natural scenes (Cerf, Paxon Frady, & Koch, 2009; Drewes, Trommershäuser, & Gegenfurtner, 2011). On each template, we manually labelled each pixel associated with a head (recall that all templates depicted animals and/or humans). The resulting masks, which covered 9% of the image area on average (SD = 12%, median = 3%), served as the ROIs for the template and its associated two-tone image. For each image and condition, we calculated the percentage of fixations landing within the ROIs (Fig. 5B). This metric increased in the After compared to the Before condition, indicating that changes in fixations were object-specific (Before: M = 30%, SD = 24; After: M = 44%, SD = 25; t(29) = 8.64, p < 0.001; M_diff_ = 0.14, 95% CI = [0.1, 0.17]). Furthermore, there were more fixations within the ROIs in the Template compared to the After condition (Template: M = 54%, SD = 25; t(29) = 6.02, p < 0.001; M_diff_ = 0.1, 95% CI = [0.06, 0.13]). Overall, the ROI analysis provides clear evidence to suggest that the influence of knowledge-dependent object representations on fixation patterns is object-specific.

### Analysis of first fixations: changes to the spatial distribution of fixations occur shortly after image onset

In order to assess the time-course of the influence of knowledge-dependent object representations on oculomotor control, we repeated our previous analyses exclusively for first fixations. This restriction did not change the overall pattern of the results (see Fig. 5C and D), suggesting that even first fixations were influenced by object representations that emerged as a consequence of the observer’s prior knowledge. Specifically, the statistical analysis showed that for first fixations, the similarity between Template and After was higher than for Template and Before (Template-After: M = 0.74, SD = 0.15; Template-Before: M = 0.62, SD = 0.17; t(29) = 4.91, p < .001; M_diff_ = 0.12, 95% CI = [0.07, 0.17]). This finding was corroborated by an ROI analysis of first fixations: the percentage of first fixations landing on ROIs was higher in the After than in the Before condition, and also higher in Template than in After (Before: M = 34%, SD = 34; After: M = 40%, SD = 35; Template: M = 60%, SD = 32; Before-After: t(29) = 3.61, p = 0.001; M_diff_ = 0.06, 95% CI = [0.03, 0.09]; Template-After: t(29) = 6.41, p < 0.001; M_diff_ = 0.2, 95% CI = [0.14, 0.27]). Taken together, these results suggest that knowledge-dependent object representations emerge fast enough to influence even the first eye-movements after stimulus onset.

### Knowledge-dependent object representations and image features act in synergy

Our analyses so far indicate that knowledge-dependent object representations play a role in gaze guidance, beginning with the first fixation after image onset. However, these analyses do not assess the role of the interaction between image-computable features and object representations. In order to address this point, we capitalized on common and distinct characteristics shared between the After condition and each of the remaining conditions (Before and Template). In particular, image-computable features of Before and After conditions are identical, but they differ in the extent to which observers experienced object representations. Specific similarities in fixation patterns between Before and the After conditions, which go beyond general factors such as centre bias, can therefore be attributed to the image-computable features of two-tone images. Conversely, the After and the Template conditions have the reverse relationship: they lead to similar object representations but differ in image-computable features. We exploited this situation to characterize the contribution of these gaze-guidance factors in the After condition.

For this purpose, we created linear combinations of heatmaps from the Before and Template conditions to compare with the heatmaps of the After condition (Fig. 6). Each new linear-combination heatmap was calculated from the Before and the Template conditions’ heatmaps, using the formula:

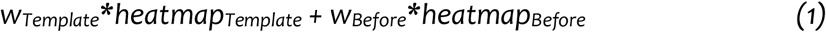

where w is a weight for the heatmap indicated by the subscript. Incorporating the normalization assumption (*w*_*Template*_ + *w*_*Before*_ = 1), we created a continuum of heatmaps spanning the range between being fully determined by the Template heatmap to being fully determined by the Before heatmap. This continuum was uniformly sampled with a step-size of 0.05. This procedure led to a set of heatmaps, which capture factors driving eye movements in the Before and the Template conditions to varying degrees. Evaluating the similarity of these new heatmaps with those from the After condition allowed us to determine the relative contribution of image-computable features and object representations to gaze guidance in the After condition. To focus on the time course, we conducted this analysis separately for first fixations and for all the remaining fixations.

**Fig. 6.**
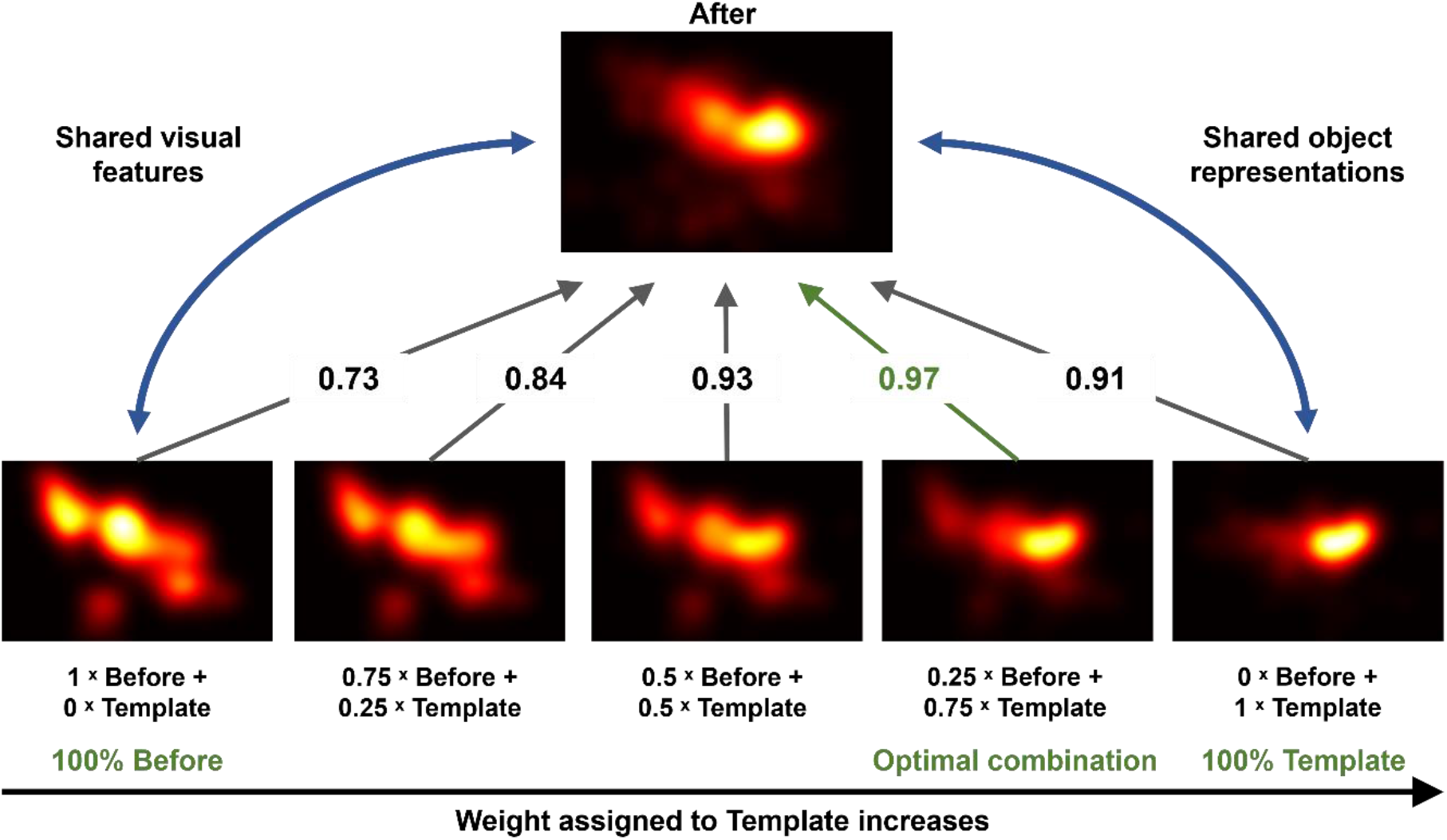
Linear combination analysis – illustration for a single two-tone image. The bottom row shows heatmaps that have been created by linearly combining the heatmaps from the Before and the Template conditions, as indicated by the text below each image. These linear-combination heatmaps were compared to the heatmap of the After condition as indicated by arrows. Numbers on the arrows indicate correlation values. The blue, double-pointed arrows illustrate the fact that the After condition shares image-computable features and object representations with the Before and the Template condition, respectively. To enable visually comparing all heatmaps shown on the figure, their pixel values were jointly normalised to zero-one range.

The results of this similarity analysis suggest that both first and all remaining fixations in the After condition were guided synergistically by image-computable features and object representations (Fig. 7). The linear-combination heatmaps that had the highest similarity with the first fixations in the After condition showed an influence from the Template heatmap but also had a substantial contribution from the Before heatmap (*w*_*Template*_ = 0.4, *w*_*Before*_ = 0.6; mean correlation M = 0.85, SD = 0.06; see Fig. 7A). Statistical analyses indicated that the heatmaps from the After condition were more similar to this optimal linear-combination heatmap than to either the Before or the Template conditions alone (Optimal-After vs. Before-After: t(29) = -2.67, p = 0.012; M_diff_ = 0.03, 95% CI = [0.01, 0.04]; Optimal-After vs. Template-After: t(29) = 5.70, p < 0.001; M_diff_ = 0.11, 95% CI = [0.07, 0.15]).

**Fig. 7.**
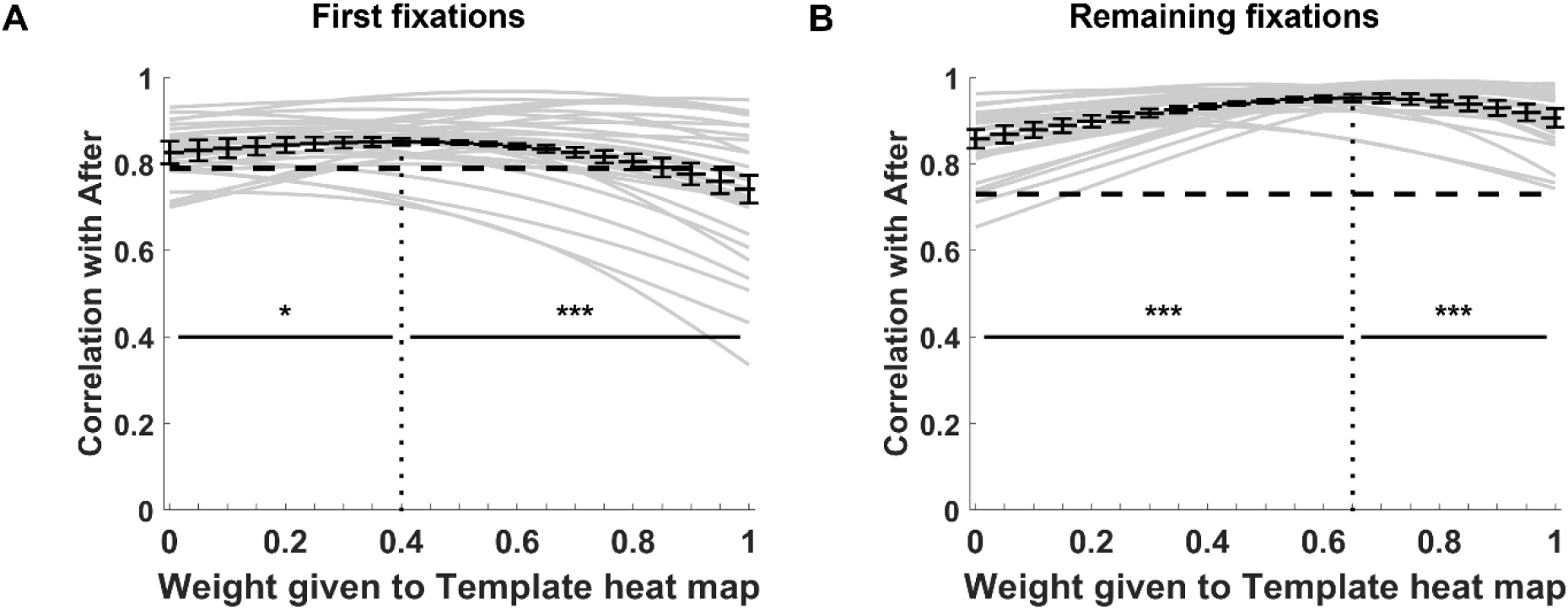
Similarities of heatmaps from the After condition to different linear combinations of heatmaps from the Template and Before conditions. A) Similarities obtained when only first fixations from the After condition are considered. B) The same analysis but for all the remaining fixations (i.e., without first) from the After condition. The weights of the linear combinations for which the similarity is maximal (indicated by the dotted vertical lines) are located more to the left for first fixations than for the remaining fixations, demonstrating that the influence of knowledge-dependent object representations increases with time. Dashed vertical lines on both panels indicate the baseline, i.e., the average similarities of the respective After heatmaps to centre bias model (M = 0.79, SD = 0.09 for first fixations; M = 0.73, SD = 0.14 for the remaining ones).

The findings for all remaining fixations from the After condition were similar (Fig. 7B). However, the linear combinations that were optimal for these fixations were more strongly influenced by the Template heatmap (*w*_*Template*_ = 0.65, *w*_*Before*_ = 35; mean correlation M = 0.95, SD = 0.03). Yet, even for these later fixations, there was a substantial influence of image-computable factors as captured by the Before heatmaps. This idea is supported by the statistical analysis, which indicates that the heatmaps from the After condition were more similar to the optimally combined heatmaps compared to both the Before and the Template condition alone (Optimal-After vs. Before-After: t(29) = 6.49, p < 0.001; M_diff_ = 0.09, 95% CI = [0.06, 0.12]; Optimal-After vs. Template-After: t(29) = 5.48, p < 0.001; M_diff_ = 0.05, 95% CI = [0.03, 0.06]).

Overall, the analysis suggests that image-computable features and object representations guide eye movements in a synergistic manner (see also Borji & Tanner, 2016). The contribution of these two factors vary over time, with object representations playing a less important role in first fixations than in later fixations. Yet, both factors already influence first fixations.

### Knowledge-dependent object representations affect multiple characteristics of oculomotor behaviour

In our final analyses of Experiment 1, we assessed the extent to which knowledge-dependent object representations affect characteristics of eye movements that might be indicative of a more fundamental change in the observers’ information-sampling strategy. First, we calculated the mean number of fixations, average fixation duration (in seconds), and average Euclidean distance between consecutive fixations (interfixation distance, in degrees of visual angle) per image, and compared them across conditions (Fig. 8). Compared to the Before condition, the After condition showed a decrease in the number of fixations (Before: M = 281.37, SD = 13.22; After: M = 240.10, SD = 19.32; t(29) = 12.76, p < 0.001; M_diff_ = 41.27, 95% CI = [34.65, 47.88]), an increase in the fixation duration (Before: M = 0.28, SD = 0.01; After: M = 0.30, SD = 0.02; t(29) = -8.22, p < 0.001; M_diff_ = -0.02, 95% CI = [0.02, 0.03]), and a decrease in interfixation distance (Before: M = 4.09, SD = 0.45; After: M = 3.34, SD = 0.55; t(29) = 11.24, p < 0.001; M_diff_ = 0.75, 95% CI = [0.61, 0.89]). We did not find statistically significant differences between the Template and the After conditions for any of these metrics (number of fixations: t(29) = -0.50, p = 0.621; M_diff_ = -2.67, 95% CI = [-13.58, 8.25]; fixation duration: t(29) = -0.24, p = 0.816; M_diff_ = 0, 95% CI = [-0.01, 0.01]; interfixation distance: t(29) = 0.32, p = 0.755; M_*diff*_ = 0.04, 95% CI = [-0.19, 0.27]; descriptive statistics for these three respective characteristics for Template condition: M = 242.77, SD = 31.76; M = 0.30, SD = 0.03; M = 3.3, SD = 0.96).

**Fig. 8.**
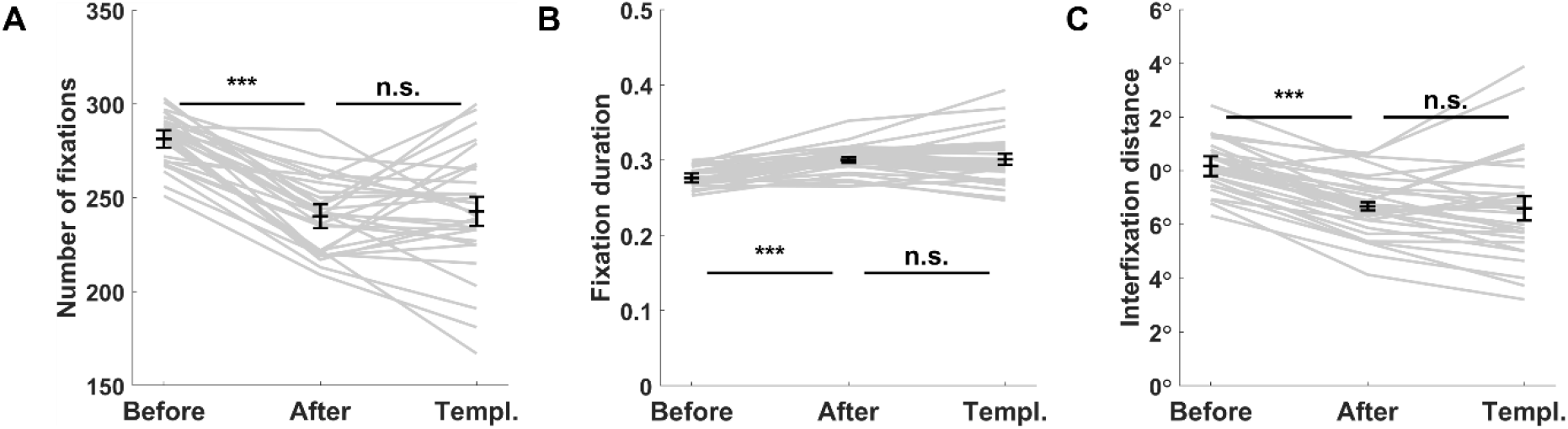
Number of fixations (A), fixation duration (B), and interfixation distance measured in degrees of a visual angle (C). All three were calculated per image and compared between conditions.

These findings are consistent with the idea that observers shift from exploring the whole stimulus in the Before condition towards extracting information only from selected parts in the After and Template conditions. To further substantiate this interpretation, we calculated the normalized entropy for the heatmaps in the different conditions (Fig. 9A). This measure is thought to index the extent to which an observer’s behaviour is exploratory (Gameiro, Kaspar, König, Nordholt, & König, 2017; Kaspar et al., 2013). Normalized entropy was lowest in the Template condition, increased in the After condition, and was highest in the Before condition (Before: M = 0.56, SD = 0.05; After: M = 0.48, SD = 0.06; Template: M = 0.42, SD = 0.07; Before-After: t(29) = 9.92, p < 0.001; M_diff_ = 0.09, 95% CI = [0.07, 0.10]; After-Template: t(29) = 6.28, p < 0.001; M_diff_ = 0.05, 95% CI = [0.04, 0.07]). In other words, observers showed the highest exploratory behaviour in the Before condition, followed by the After and the Template condition.

**Fig. 9.**
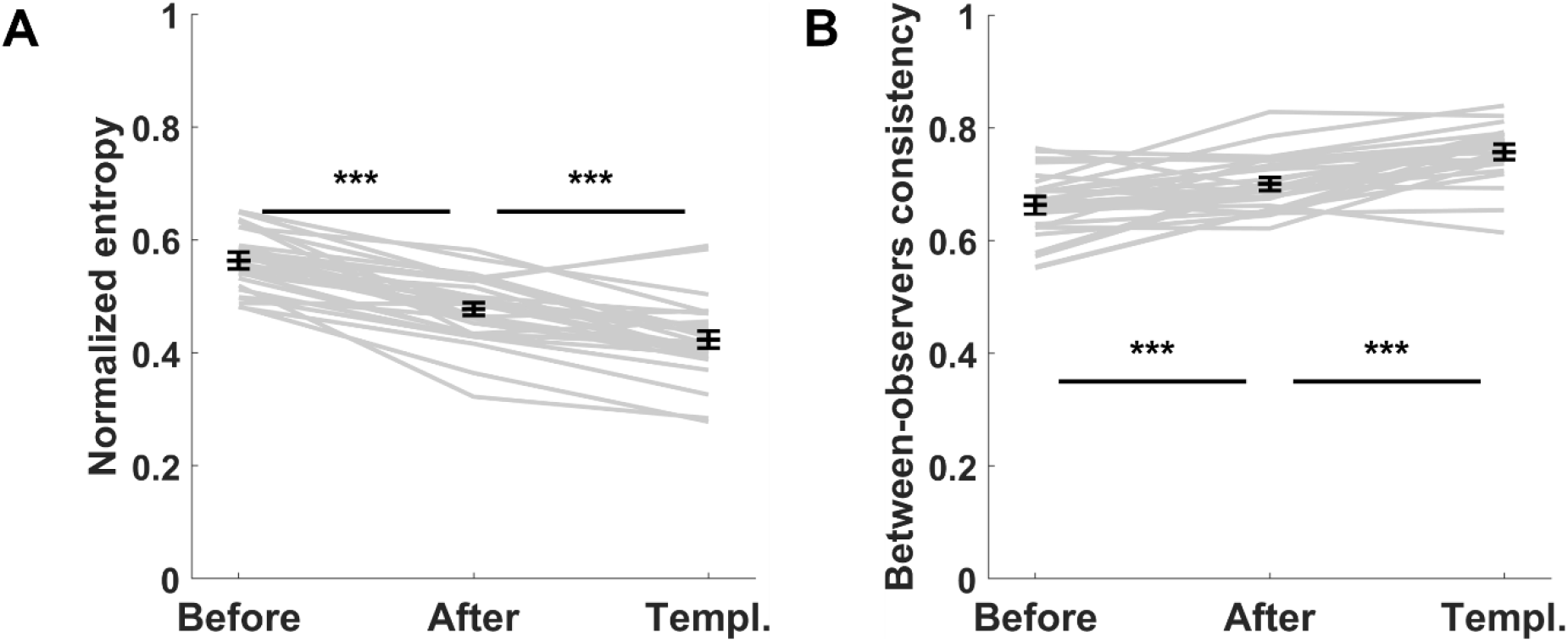
Normalized entropy and between-observers consistency. A) Normalized entropy of fixation distributions (in arbitrary units) as a measure of their spread. Higher values indicate more exploratory behaviour of observers. B) Between-observers consistency in selecting fixation targets measured by how similar (on average) fixations of a single observer were to fixations of all the remaining observers pooled together.

In our final analysis, we wanted to know if object representations would result in more homogenous gaze behaviour across observers (Fig. 9B). We quantified between-observers consistency by averaging the similarity between each observer’s individual heatmap to the heatmaps of all remaining observers (Lyu et al., 2020). This metric increased both between the Before and After conditions and between the After and Template conditions (Before: M = 0.66, SD = 0.05; After: M = 0.7, SD = 0.05; Template: M = 0.76, SD = 0.05; Before-After: t(29) = 3.96, p < 0.001; M_diff_ = 0.04, 95% CI = [0.02, 0.06]; After-Template t(29) = 6.96, p < 0.001; M_diff_ = 0.06, 95% CI = [0.04, 0.07]), suggesting that object representations increase consistency in information-sampling behaviour across observers.

## Experiment 1 – Discussion

In Experiment 1, we measured eye-movements in response to grayscale images of scenes containing objects and two-tone images derived from these templates. On initial viewing, two-tone images are experienced as meaningless black-and-white patches. Once an observer has acquired relevant prior object-knowledge, however, the visual system organizes the patches into a coherent percept of an object. We demonstrate that, when a two-tone image is perceived as showing a coherent object rather than meaningless patches, gaze guidance changes in several ways. First, and most importantly, fixation patterns on two-tone images become more similar to those measured in response to the template when two-tones lead to object representations vs. when they are experienced as meaningless patches. Moreover, fixation locations become more object-specific. Importantly, however, we also demonstrate that object representations do not fully dominate gaze guidance, but that image-computable feature space and object representations interact in determining where people look. While the data suggest a specific temporal development of this interaction, we also observe that the influence of knowledge-dependent object representations is already present in the first eye-movement after image onset. Object representations also lead to fewer fixations, longer fixation durations, shorter interfixation distances as well as a less exploratory pattern of eye movements and more consistency across observers. Overall, these results suggest that object representations, which are not fully determined by image-computable features but depend on an observer’s prior object-knowledge have a substantial influence on eye movements.

It is, however, possible that the change in fixation patterns observed in Experiment 1 were caused by a memory process unrelated to knowledge-driven perceptual organization. Specifically, it has been suggested that eye movements preformed during memory retrieval of an image resemble the eye movements performed when seeing this stimulus for the first time (Noton & Stark, 1971; see Wynn, Shen, & Ryan, 2019 for a recent review and Foulsham & Kingstone, 2013 for criticism). According to this alternative explanation, two-tone images in the After condition might have acted as cues that triggered the retrieval of the corresponding template, and this retrieval might have been accompanied by the re-enactment of gaze behaviour from the Template condition. A simpler but overall similar alternative explanation of the results from Experiment 1 might suggest that memory-retrieval of template images resulted in the observers voluntarily directing their gaze towards locations in the two-tone images, which they remembered to be occupied by objects. According to both explanations, the factor driving changes in eye movements in the After condition is the mapping of objects to locations that the observers remember from the Template condition, rather than perceptual organization induced by prior object-knowledge. To exclude these alternative interpretations, which we label ‘object-to-location mapping hypothesis’, we conducted Experiment 2.

### Experiment 2

Experiment 2 was identical to Experiment 1 in all aspect except that the template images were flipped along the vertical axis (‘mirror-flipped’) from left to right. Consequently, the screen locations occupied by objects differed between the Template condition and the remaining conditions. This simple manipulation allowed us to adjudicate between the different alternative interpretations mentioned in the previous section: according to the object-to-location mapping hypothesis, which suggests that observers merely revisited the parts of the display, which contained objects during the presentation of template images, we would expect a high similarity between heatmaps from the After and Template conditions, despite the lack of overlap in spatial location of objects in these two conditions. If, however, the effects observed in Experiment 1 were attributable to knowledge-dependent object representations, we would expect the similarity between the After and Template conditions to be low (see Fig. 3 for illustration). Moreover, by mirror-flipping the heatmaps obtained from the mirror-flipped templates, we would expect an increase in similarity to levels seen in Experiment 1 (because this leads to a re-alignment of heatmaps from templates and two-tones).

### Experiment 2 – Method

A separate set of 18 Cardiff University students (mean age 19.5 years, 5 males), who did not participate in Experiment 1, served as observers. The design of Experiment 2 was identical to that of Experiment 1 except that the template images were flipped along the vertical axis from left to right during all parts of the experiment (i.e., during instructions, the Blending Phase, and the Template condition). This condition is labelled FlippedTemplate.

## Experiment 2 – Results and Discussion

### Memory-retrieval of object-to-location mapping does not explain changes in eye movements

Similar to Experiment 1, the meaningfulness ratings provided by the observers after viewing each two-tone were higher in the After condition than the Before condition both when we averaged them per observer (t(17) = 6.62, p < 0.001; M_diff_ = 0.24, 95% CI = [0.16, 0.31]) and per image (t(29) = 16.74, p < 0.001; M_diff_ = 0.24, 95% CI = [0.21, 0.27]). This result indicates that observers were able to bind the two-tone images into meaningful percepts despite viewing templates, which were presented in a mirror-flipped manner.

The results of the eye-movements data analysis were inconsistent with the object-to-location hypothesis but provided support for the idea that knowledge-dependent object representations influence eye movements (see Fig. 10). In particular, by contrast to the analogous analysis in Experiment 1, heatmap similarities did not differ when comparing the FlippedTemplate-Before pair vs. the FlippedTemplate-After pair (FlippedTemplate-Before: M = 0.46, SD = 0.22; FlippedTemplate-After: M = 0.48, SD = 0.22; t(29) = 1.45, p = 0.158; M_diff_ = 0.03, 95% CI = [-0.01, 0.06]). A BF of 0.50 suggested that the data provided evidence in favour of there being no difference between conditions, but that this evidence was weak. Importantly, once the heatmaps from the template and two-tone images were re-aligned, by flipping the heatmaps of the FlippedTemplate condition, the similarity between the RealignedTemplate and the After condition was higher than the similarity between RealignedTemplate and Before (RealignedTemplate-Before: M = 0.68, SD = 0.15; RealignedTemplate-After M = 0.8, SD = 0.11; t(29) = 7.77, p < 0.001; M_diff_ = 0.13, 95% CI = [0.09, 0.16]).

**Fig. 10.**
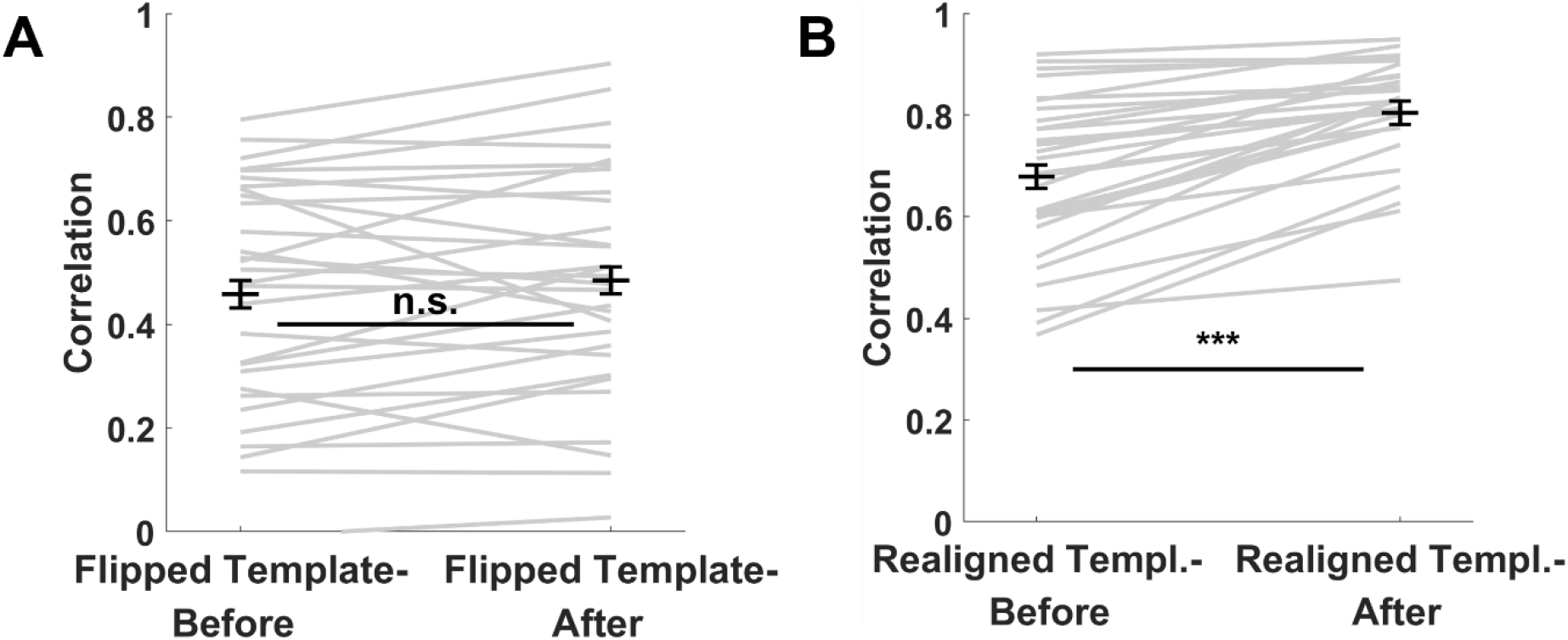
Results of Experiment 2. Similarities between heatmaps from two-tone images and mirror-flipped templates, where the two-tones were viewed either in the Before or in the After condition. The heatmaps derived from the mirror-flipped template images were used either before (A) or after (B) the mirror-flipping was reverted by ‘flipping back’ these heatmaps and realigning them with the heatmaps from two-tone images.

In sum, the results from Experiment 2 exclude the possibility that gaze guidance in the After condition is based on a mapping of objects to locations via retrieval of this information from the Template condition. In a third experiment, we addressed two further alternative explanations of the results from Experiment 1. First, it is possible that during the phase when two-tone images are blended with templates, observers learn to associate specific image-features in the two-tone images with objects from the templates. When viewing two-tone images in the After condition, these feature-object associations might guide fixations towards these specific visual patterns, irrespective of transformations such as those introduced by the mirror-flipping. While this possibility might seem implausible, there is evidence to suggest that such learning processes are an important factor in oculomotor control (Alfandari, Belopolsky, & Olivers, 2019).

A final alternative explanation of our results from both Experiment 1 and 2 relates to potential order effects. It is possible that the changes in fixation patterns between Before and After conditions resulted from viewing two-tones for a second time, rather than from knowledge-dependent perceptual organization. In other words, observers might sample information from different image regions on second compared to first viewing, irrespective of the kind of information they acquire in the meantime. We conducted Experiment 3 to exclude the possibility that (i) feature-object associations, or (ii) any order effects could explain the effects of Experiments 1 and 2.

### Experiment 3

Experiment 3 adopted the same procedure as the previous experiments except that the templates from Experiment 1 (‘real templates’) were replaced with different images that were unrelated to the two-tones (‘dummy templates’). This experimental design allowed us to test whether feature-object associations provide a plausible explanation for the findings of Experiment 1 and 2. Specifically, observers might associate certain features in the two-tone images with objects in the templates during the Blending Phase. When viewing two-tone images in the After condition, these feature-object associations could drive fixations towards image locations in the two-tones that overlap with objects in the respective (dummy) templates. These effects should be observable despite observers not having acquired the prior object-knowledge required to organize the two-tone images into coherent percepts. Moreover, the design also allowed us to assess whether order effects could explain the findings from Experiments 1 and 2.

### Experiment 3 – Method

Experiment 3 was completed by 20 observers (mean age 19.55, 5 males) who did not participate in the previous two experiments. All were Cardiff University students. The procedure was identical to the previous experiments except that in each block, the templates used in the Template condition and in the Blending Phase were unrelated to the two-tones presented in this block (‘dummy templates’). Each two-tone had a unique dummy template paired with it and this pairing was fixed for all observers. Importantly, each dummy template was a ‘real template’ of a different two-tone presented in the preceding block during the experiment (see Fig. 11). While templates in this experiment could thus not provide object knowledge that would help organize the two-tone image into an object percept in the After condition, we were nevertheless able to register eye movements on the real templates. Measuring fixations on real templates was necessary to assess whether simply viewing a two-tone for a second time, without prior object-knowledge, would lead to increased similarity between heatmaps of two-tone images in the After and their real templates, as seen in the previous experiments.

**Fig. 11.**
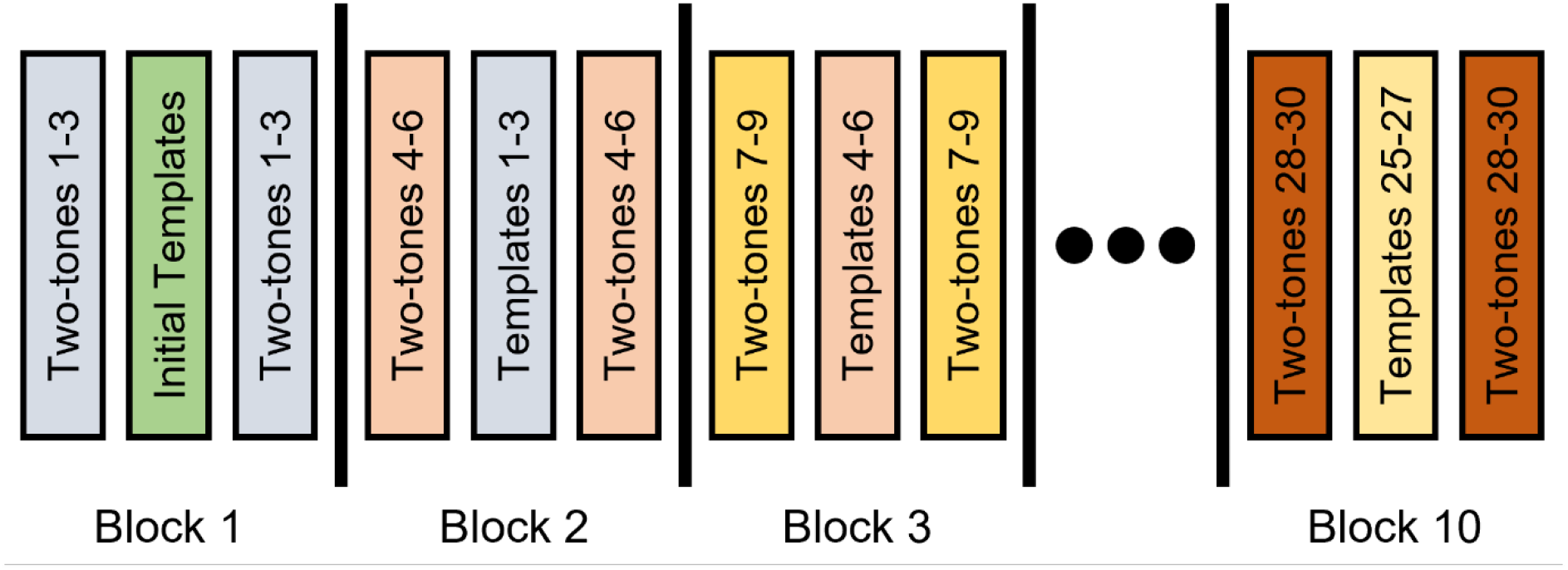
Randomization schema used in Experiment 3. Within each block, stimuli were presented in a randomised order (as in Experiment 1 and 2). The presentation of images was arranged in such a way that templates in, e.g., Block 2, were the real templates of the two-tone images in Block 1. This order allowed us to register fixations for the real templates (for comparison with fixation on two-tone images) while omitting the opportunity for the observer to acquire the relevant prior object-knowledge that would allow them to disambiguate the two-tone images.

In the first block, the same dummy templates – greyscale images not related to any of the two-tones – were always presented. In all other blocks, the assignment of stimuli to experimental blocks was pseudo-randomized for each observer individually in a way which guaranteed that dummy templates presented in any given block were the real templates of two-tones presented in the preceding block (see Fig. 11). To ensure that we included data from the same number of observers for each two-tone and template, we had to discard fixations registered on the two-tones presented in the final experimental block and fixations from the dummy templates from the first blocks (‘initial templates’). Note that – because we pseudo-randomized the order of stimulus presentation for each observer individually – for different images we had to discard data from different observers. Importantly, however, for each image set consisting of a two-tone (viewed in Before and After condition), its dummy template, and its real template, we retained data from a homogenous group of 18 observers (out of 20 who completed the experiment), but the composition of these groups was different for different image sets.

## Experiment 3 – Results

### Lack of relevant object-knowledge prevents the emergence of knowledge-dependent object representations

The analysis of meaningfulness ratings demonstrated that, as expected, observers were not able to bind the two-tone images into coherent object percepts even in the After condition (Fig. 12A and B). In particular, the differences in ratings between Before and After conditions were not statistically significant, both when the data were averaged per observer (t(19) = 1.49, p = 0.152; M_diff_ = 0.02, 95% CI = [-0.01, 0.06]) or per image (t(29) = 1.97, p = 0.058; M_diff_ = 0.02, 95% CI = [0, 0.05]). In the former case, Bayes factor analysis suggested weak evidence for the lack of differences (BF = 0.60), while in the latter no clear conclusions could be drawn (BF = 1.07).

**Fig. 12.**
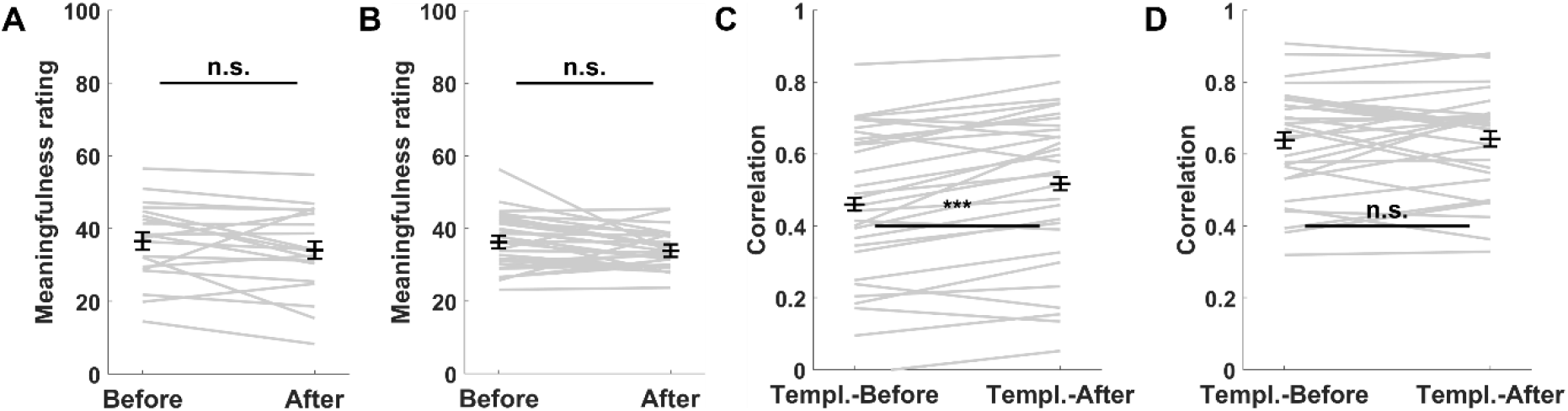
Results of Experiment 3. Meaningfulness ratings averaged per observer (A) and per image (B). As expected, the meaningfulness ratings suggest that observers did not perceive the two-tones in the After condition as more meaningful compared to the Before condition. C) Comparison of heatmap similarities between two-tones (viewed in the Before and After conditions) and their dummy templates (i.e., unrelated images). This comparison revealed that, in the absence of the relevant prior object-knowledge, observers’ eye-movements in the After condition were slightly influenced by the memory of dummy templates but this effect was too small to explain the effect seen in Experiment 1. D) Comparison of heatmap similarities between two-tones (viewed in Before and After conditions) and their real templates. Viewing the two-tone images for a second time without having seen the real templates does not result in gaze patterns that are more similar to the ones recorded on their real templates compared to first viewings.

### Memory-retrieval of feature-object associations leads to small changes in eye movements but cannot explain key findings of Experiment 1

Experiment 3 tested the hypothesis that the effects observed in the two previous experiments might be explainable by a learned association between feature clusters in two-tones and object locations on templates. Specifically, it is possible that during blending of two-tone images and templates, observers learn to associate specific features of the two-tones with object locations in the templates and then re-visit these features when viewing the two-tone images in the After condition. Our analysis indicated that the similarity in heatmaps in the DummyTemplate-After pair was higher compared to the DummyTemplate-Before pair (Fig. 12C). This increase in similarity, although significant in a statistical sense, was small (DummyTemplate-Before: M = 0.46, SD = 0.21; DummyTemplate-After: M = 0.52, SD = 0.22; t(29) = 4.70, p < 0.001; M_diff_ = 0.06, 95% CI = [0.03, 0.08]). Nevertheless, the analysis provided evidence to suggest that feature-object associations might guide oculomotor control to a limited extent. Alternatively, these results could be driven by memory retrieval of object-locations in the templates: while Experiment 2 showed that memory retrieval does not play a role when perceptual organization takes place, this process may become important when the stimulus remains unorganized with no object representations to guide eye movements.

While these results suggest that object-to-location or object-to-feature mapping might influence gaze guidance in the After condition, the key question is whether these effects can explain the results found in Experiment 1. To address this issue, we directly compared the increase in similarity between the Template-Before vs. Template-After pairs across Experiments 1 and 3. Given that both experiments differed with respect to the number of observers who contributed to the heatmaps of each image, we included fixations only from 18 observers from Experiment 1 (drawn randomly). To ensure that the outcome does not depend on the specific set of observers from Experiment 1, we repeated this analysis for 20 different, randomly drawn sets and obtained the same pattern of outcomes for each of them. Specifically, the analysis indicates that the change in similarity between the Template-Before vs. Template-After pairs was larger in Experiment 1 than in Experiment 3 (Experiment 1: M = 0.17, SD = 0.13; Experiment 3: M = 0.06, SD = 0.07; t(29) = 4.15, p < 0.001; M_diff_ = 0.11, 95% CI = [0.06, 0.17]). Our results thus demonstrate that the processes responsible for changing gaze-patterns between Before and After conditions in Experiment 3 cannot fully explain the analogous changes in Experiment 1.

### The findings of Experiment 1 cannot be attributed to order effects

In the final analysis, we considered the possibility that order effects explain the key findings of Experiment 1 and 2. Specifically, we asked whether viewing the same two-tones for a second time without receiving prior object-knowledge could change fixation patterns such that they would resemble the patterns from the (real) templates. Recall that the design of Experiment 3 ensured that observers saw each two-tone image twice, each time without prior object-knowledge (Before and After conditions, respectively) and they also saw the real template for these two-tones in the following block. If the findings in Experiments 1 and 2 resulted, at least partly, from an order effect, we would expect that the similarity in fixation patterns in the (real) Template-After pair would be higher than in the (real) Template-Before pair in the current experiment.

The results were inconsistent with this ‘second-viewing’ hypothesis (Fig. 12D). The heatmap similarities between the real templates and the corresponding two-tones viewed in the Before and After conditions were not statistically different (Template-Before M = 0.64, SD = 0.15; Template-After M = 0.64, SD = 0.14; t(29) = 0.22, p = 0.830; M_diff_ = 0, 95% CI = [-0.03, 0.03]). Moreover, a Bayes factor analysis provided evidence to support a lack of a difference (BF = 0.20).

## Discussion

When an observer explores the environment with no specific task other than to obtain information, control of eye movements is typically considered within a dichotomy between low-level features and high-level object representations. Here, we abandoned this simplifying framework in light of emerging evidence highlighting the complex and intricate relationship between image-computable features and high-level object representations in visual perception. We recorded eye movements in response to two-tone images, stimuli that appear as meaningless patches on initial viewing but, once relevant object-knowledge has been acquired, are organized into coherent and meaningful percepts of objects. In the current study, prior object-knowledge was provided in the form of template images, i.e., the unambiguous photographs from which the two-tone images had been generated. Across three experiments, fixation patterns on the same two-tone images differed substantially depending on whether observers experienced them as meaningless patches or organized them into object representations. In particular, when organized into object representations, we found that fixation patterns on two-tone images were more similar to those on templates, more focused on object-specific, pre-defined regions-of-interest, less dispersed, and more consistent across observers. These effects were evident from the first fixations on an image. Importantly, eye-movements on two-tone images were best explained by a simple computational model that takes into account both low-level features and high-level, knowledge-dependent object representations. Together, these findings highlight the importance of dynamic interactions between image-computable features and knowledge-driven perceptual organization in guiding information sampling via eye-movements in humans.

The idea that knowledge-driven object representations restructure human eye-movements is supported by both our general assessment of fixation distributions between two-tone images and template, and also by a more specific analysis focusing on fixations within regions-of-interest. These findings provide strong support for the hypothesis that objecthood *per se* contributes to the process of selecting fixation targets in images. In our experimental design, image-computable visual features are insufficient for object representations to emerge, their formation is dependent on prior object-knowledge. On the one hand, this characteristic of two-tone images is an important experimental tool: it allows us to decisively rule out the possibility that human oculomotor control during free viewing relies solely on image-computable features, regardless of whether these features are low- or high-level (Zelinsky & Bisley, 2015). The simple but critical result in this regard is the finding that eye-movement patterns differed dependent on whether observers had formed object representations despite the fact that the features in the stimuli remained identical. In addition to its use as an experimental tool, however, the dependence of object representations on prior knowledge is also important from a conceptual perspective. Specifically, the finding that fixations were guided by knowledge-dependent representations demonstrates that for the oculomotor system, objects cannot be conceptualised (exclusively) as image-computable, high-level features in the way many previous studies have done (Borji & Tanner, 2016; Einhäuser et al., 2008; Schütt et al., 2019; Stoll et al., 2015). Rather, objecthood that is relevant for guiding eye-movements is a characteristic that is distinct from the collection of any low- or high-level features. In our study, objecthood emerges in the interaction between prior object-knowledge and the visual input. Whether object representations that are relevant for oculomotor control are always distinct from the featural input is a difficult question that we cannot answer with our data. However, the size, the speed, and the incidental nature of these effects suggests that they might be characteristic of eye-movement control in everyday visual behaviour.

Equally important as the finding that knowledge-driven object representations guide human gaze is the fact that they do not fully determine the selection of fixation locations. While eye-movements on two-tone images changed once they elicited object representations such that fixation distributions became more similar to fixations on template images, substantial differences in eye-movements remained between these two conditions. Our linear combination analysis suggests that this disparity is systematic and can be explained by the differences in the features in two-tone vs. template images. In this analysis, we generated linear combinations of the fixation patterns registered on template images and on two-tone images that did not elicit object representations (i.e., Template and Before conditions, respectively). Specifically, these linear combinations included varying proportions of the heatmaps of both viewing conditions. We then assessed the similarities between the combined heatmaps and the heatmaps that resulted from the condition, in which the two-tone images elicited object representations (After condition). These similarities peaked at a point at which the combined heatmaps were determined by the fixation distributions from both the Template and the Before conditions. The finding thus demonstrates that when observers experienced the percept of an object in the two-tone images (After condition), fixations were best explained by a combination of the factors guiding eye movements in the Before and the Template conditions. Specifically, even when observers perceived an object in the two-tone images, their eye movements were only partly determined by the factors that guide eye movements in response to the template image. The image-computable features that drive eye movements in response to two-tone images when no object is perceived (Before condition) still made a substantial contribution to gaze guidance. Note that the linear combination analysis was conducted on a per-image basis. The finding that both features and objecthood contribute to eye-movement control can therefore not be explained by averaging across different images, with some leading to purely feature-driven and other to purely representation-driven eye-movement control.

The finding that features remain important for eye-movement control even after having been bound into a high-level object representation potentially challenges some of the strong claims regarding the role of features vs. objects in gaze guidance. For instance, the cognitive relevance theory (Henderson et al., 2009) proposes that visual features do not contribute to oculomotor control directly but provide the means to generate a representation of potential fixation locations that have not yet been ranked for priority. High-level factors operate on this ‘flat landscape’ to determine the ultimate fixation locations. In other words, features are important only as potential carriers of higher-level representations and do not contribute to eye-movement control by themselves. According to this idea, it follows that as long as visual features give rise to similar object representation, these representations should guide eye movements towards similar locations. Therefore, to the extent to which two-tones and templates lead to similar object representations, we would have expected both image types to result in similar eye-movement patterns independent of their featural differences. Contrasting with this notion, we found that, while features can be flexible carriers of object representations that guide eye-movements as predicted by the cognitive relevance theory, the specific features that support these high-level representations persist to exert a sizeable influence.

In terms of the time-course of eye-movements, we provide clear evidence that already the first fixations after image onset are affected by objecthood. Interestingly, however, the linear combination analysis indicates that for first fixations the relative influence of features is stronger – and, therefore, the relative influence of objecthood weaker – compared to later fixations. Thus, while the influence of knowledge-dependent object representations emerges quickly, the linear combination analysis suggests that the effects of knowledge-driven perceptual organization continue to build beyond the first fixation, by contrast to the effects of features. Nevertheless, our data suggest that the influence of knowledge-dependent object representations emerges quickly and exerts an influence from the earliest fixations.

At image onset, when the eyes are stationary prior to the first saccade, most of the image is viewed via peripheral vision with only a small part being inspected with high-resolution foveal vision. The analysis of the first fixations therefore suggest that the visual system is able to generate knowledge-dependent object representations quickly and largely based on information from peripheral vision. Due to the optical, anatomical, and neurophysiological characteristics of the primate visual system, peripheral vision is limited in various respects (Rosenholtz, 2016), but there is good evidence that it provides enough information to generate a gist representation of a visual scene that can guide subsequent eye movements (Anderson, Donk, & Meeter, 2016; Castelhano & Henderson, 2007; Melissa L.H. Võ & Schneider, 2010). Exactly how detailed this gist representation is, which features it contains, and whether objects are represented varies depending on a number of different factors (Crouzet, Joubert, Thorpe, & Fabre-Thorpe, 2012; Groen et al., 2013; Larson & Loschky, 2009; Malcolm, Groen, & Baker, 2016; Wallis, Bethge, & Wichmann, 2016). Note, however, that this question is of limited relevance in the current context because features in two-tone images – independently of whether they are viewed by foveal or peripheral vision – are necessary but, by themselves, not sufficient to determine the high-level object representations we study here. However, one notion that might help in explaining the rapid influence of knowledge-dependent object representations on eye movements is provided by the suggestion that object recognition involves a predictive process that is triggered by low spatial-frequencies in the input (Bar et al., 2006; Bar, 2003, 2004, 2021; Bullier 2001). Specifically, low spatial-frequency information is thought to be fed forward by fast projections to high-level brain systems that connects this rudimentary input to prior object-knowledge. This process narrows down the search space of possible hypotheses about object identities in the input, thereby scaffolding and shaping a more precise perceptual experience of the input. It is therefore tempting to speculate that, in our experiment, first fixations were guided by object representations that are based on the process that links impoverished low spatial-frequency image content to prior knowledge, while later fixations might be based on fuller object representations. This idea rests on the assumption that two-tone images provide low spatial-frequency information to peripheral vision that allows the linking of two-tone images to memory representations of template images. Given that the image-processing operations required to generate two-tone images mainly affect high spatial-frequency components and have less impact on low spatial frequencies, this assumption seems plausible.

While our analyses mainly focused on locations of fixations, other aspects of oculomotor control are also influenced by knowledge-dependent perceptual organization. Specifically, we observed a decrease in saccade length and an increase in fixation duration when two-tone images were organized into object representations (After condition) compared to when they were not (Before condition). Both changes are indicative of a shift from image exploration to image exploitation (Gameiro et al., 2017; Kaspar et al., 2013), an interpretation that was also supported by the decrease in entropy across the two conditions. The oculomotor system constantly has to decide whether to keep the eyes still in order to be able to further inspect the currently fixated scene region – a process referred to as exploitation –, or to perform a saccade to explore another part of the image. Interestingly, in our study, the shift from exploration to exploitation went along with an increase in the amount of fixations landing on objects. This finding suggests that the visual system prioritizes objects in a specific way: it exploits object locations for further information while abandoning exploration of the remaining parts of the image. In other words, our data demonstrate that clusters of features that are bound into, and provide support for, object representations become interesting for the visual system over non-object related feature clusters (for a similar finding, see Król & Król, 2019). The shift from exploitation to exploration once objecthood is established also leads to higher consistency across observers. This finding suggests that guidance of exploration is either more idiosyncratic or that image-computable features that are not bound into object representations do not provide strong constraints for oculomotor control. Conversely, object representations, even when supported by exactly the same features, have a structuring or normative effect on information sampling. In other words, while observers explore features in different ways, they exploit objects in similar ways.

In summary, we demonstrate that gaze guidance is best understood by dynamic interactions between image-computable features and knowledge-dependent perceptual organization. Specifically, our findings demonstrate the importance of objecthood *per se* – i.e., representations that are not reducible to image-computable features – in oculomotor control but also indicate a persistent contribution of object-independent features. We demonstrate that when visual input remains identical, the emergence of knowledge-dependent object representations substantially restructures information sampling via eye-movements. However, we also show that even when image-computable features are bound into object representations, they still retain some influence on eye movements, challenging the idea that the role of features is limited to being carriers for high-level representation without direct influence on eye-movements. Finally, we also show that the emergence of object representations results in an overall change of the information-sampling strategy of the visual system, leading to the prioritization of information extraction from features that are bound into object representations, at the expense of exploration of the entire image.

## CRediT authorship statement

M.P.: Conceptualisation, Methodology, Software, Validation, Formal analysis, Investigation, Visualization, Writing - Original Draft, Writing - Review & Editing

E. v.d. H.: Methodology, Writing - Review & Editing

C.T.: Conceptualisation, Methodology, Writing - Original Draft, Writing - Review & Editing, Supervision, Resources

