## supplement for "Knowledge-driven perceptual organization reshapes information sampling via eye movements"

**Comparison of gaze-patterns between in After and Before conditions**

As described in the main text, we find higher similarities between the heatmaps of the Template-After compared to the Template-Before pair, suggesting that knowledge-dependent object representations contribute to gaze guidance. Another prediction that can be derived from this notion relates to the heatmaps from two-tone images alone. Specifically, for a given two-tone image, heatmaps generated from fixations of two different groups of observers should be more similar to each other when each group viewed the image in the same condition (Before-Before or After-After), as compared to when the conditions were different (Before-After).

In the current analysis, we tested this hypothesis. We randomly split our sample of 36 observers into two equally large groups, and for each group generated heatmaps for the Before and After conditions. Next, for all possible pairs of conditions (Before-Before, After-After, and Before-After), we calculated similarities between heatmaps originating from different groups. If object representations affect oculomotor control, then the similarity for the Before-After pairs should be lower than the similarity for the Before-Before and the After-After pairs. The results confirmed these expectations (see Fig. S1). The similarity between heatmaps from the Before-After pairs was lower than the similarities from the Before-Before and the After-After pairs (Before-Before: $M = 0.94$, $SD = 0.03$; Before-After: $M = 0.84$, $SD = 0.09$; After-After: $M = 0.95$, $SD = 0.03$; Before-Before vs. Before-After: $t(29) = 5.94$, $p < 0.001$; $M_{diff} = 0.1$, 95% CI = [0.06, 0.13]; After-After vs. Before-After: $t(29) = 7.14$, $p < 0.001$; $M_{diff} = 0.11$, 95% CI = [0.08, 0.14]). This finding provides further evidence for the influence of knowledge-dependent object representations in oculomotor control. Note that while we compared heatmaps across groups of observers, we relied on paired sample t-tests because individual images remain the unit of analysis. In order to warrant that the outcome of this analysis did not depend on a specific composition of the two groups, we repeated the split 20 times, each time assigning observers to the groups randomly. For each spilt, we obtained the same pattern of results.

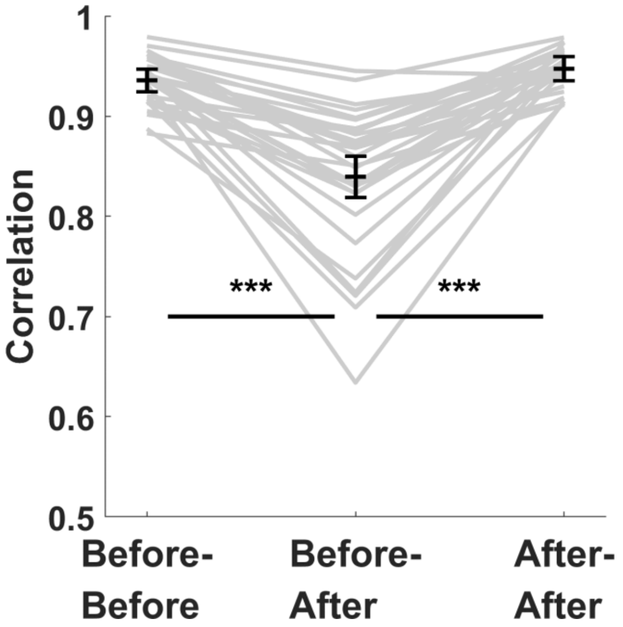


*Fig. S1. Similarities between heatmaps generated after splitting the sample from Experiment 1 into two groups, compared between conditions. As predicted, viewing two-tone images in a different condition be each group (Before-After) resulted in lower similarities than viewing in the same condition (Before-Before and After-After). Note that y-axis on the blot starts at 0.5.*

**Key findings of Experiment 1 do not depend on a specific similarity metric**

All comparisons of gaze patterns reported in the main text rely exclusively on correlation, which is one of many metrics to quantify the similarity between two heatmaps (Bylinskii, Judd, Oliva, Torralba, & Durand, 2016). To ensure that our conclusions do not depend on this specific metric, we repeated key analyses from the main text using another metric, SIM, as implemented by Bylinskii and colleagues (Kümmerer et al., 2020). SIM scores can be interpreted in a similar fashion as correlation: zero indicates no overlap between two smooth distributions, one indicates that they are identical.

First, using SIM instead of correlation, we re-run the analysis of Experiment 1 reported in the section *Knowledge-dependent object representations control the spatial distributions of fixations*. Again, heatmaps in the After condition, as compared to the Before, were more similar to heatmaps from the Template, both for all fixations (Before: M = 0.62, SD = 0.08;

After: $M = 0.76$, $SD = 0.06$; $t(29) = 10.05$, $p < .001$; $M_{diff} = 0.14$, 95% CI = [0.11, 0.17]) and for first fixations (Before: $M = 0.54$, $SD = 0.11$; After: $M = 0.61$, $SD = 0.10$; $t(29) = 4.06$, $p < .001$; $M_{diff} = 0.07$, 95% CI = [0.04, 0.11]).

Second, we re-run the analysis from the section *Knowledge-dependent object representations and image features act in synergy*, which describes the linear-combination analysis of the data from Experiment 1. We found the same pattern of results for SIM as we did for correlation. Similarly as for correlation, the heatmaps resulting from the optimal weighting for both first and remaining fixations (mean SIM value for first fixations: $M = 0.65$, $SD = 0.07$; for remaining fixations: $M = 0.84$, $SD = 0.04$) were more similar to the heatmap from the After condition than those resulting from either the Before or Template conditions alone (First: Optimal-After vs. Before-After: $t(29) = 2.61$, $p = .014$; $M_{diff} = 0.03$, 95% CI = [0.01, 0.05]; Optimal-After vs. Template-After: $t(29) = 3.16$, $p = .004$; $M_{diff} = 0.03$, 95% CI = [0.01, 0.05]; Remaining: (Optimal-After vs. Before-After: $t(29) = 8.23$, $p < .001$; $M_{diff} = 0.09$, 95% CI = [0.07, 0.12]; Optimal-After vs. Template-After: $t(29) = 7.68$, $p < .001$; $M_{diff} = 0.08$, 95% CI = [0.06, 0.10]). Moreover, replicating the pattern from the analysis reported in the main text, the optimal weighting for first fixations was closer (albeit here only slightly) to the Before heatmaps and, consequently, further from Template heatmaps than for all remaining fixations (First: $w_{Template} = 0.65$; Remaining: vs. $w_{Template} = 0.70$).

**Linear-combination analysis results differ between Experiments 1 and 3**

Experiment 3 provides an interesting opportunity to test the validity of our linear-combination analysis in Experiment 1. Recall that the experimental design of Experiment 3 guaranteed that observers did not receive prior object-knowledge between the Before and the After condition and should therefore be unable to perceptually organized the two-tone images in both conditions. Contrasting with the findings in Experiment 1, we would therefore expect that the linear-combination analysis would indicate an optimal weighting of Before and Template heatmaps that is very close, or identical to the Before heatmap on its own.

We found that the optimal weighting is the same for first and remaining fixations and amounts to 0.15 (mean correlation for first fixations: M = 0.80, SD = 0.08; for remaining fixations: M = 0.91, SD = 0.05; see Fig. S2). For the first fixations, the Optimal-After similarities did not differ statistically from the Before-After ($t(29) = 1.69$, $p = 0.101$; $M_{diff} = 0$, 95% CI = [-0.01, 0]). For the Optimal-After vs. Template-After, the difference was statistically significant ($t(29) = 9.17$, $p < 0.001$; $M_{diff} = 0.35$, 95% CI = [0.28, 0.43]). For the remaining fixations, both comparisons yielded statistically significant results (Optimal-After vs. Before-After: $t(29) = 3.61$, $p = 0.001$; $M_{diff} = 0.01$, 95% CI = [0, 0.01]; Optimal-After vs. Template-After: $t(29) = 11.56$, $p < 0.001$; $M_{diff} = 0.4$, 95% CI = [0.33, 0.47]).

Although for first as well as remaining fixations the optimal linear-combination still included both Before and Template heatmaps, in both cases the Optimal-After similarities were only marginally different from the Before-After (the 'edge' of the spectrum of linear combinations) similarities, and markedly different from the Template-After (the other edge), which was expected, given the design of Experiment 3.

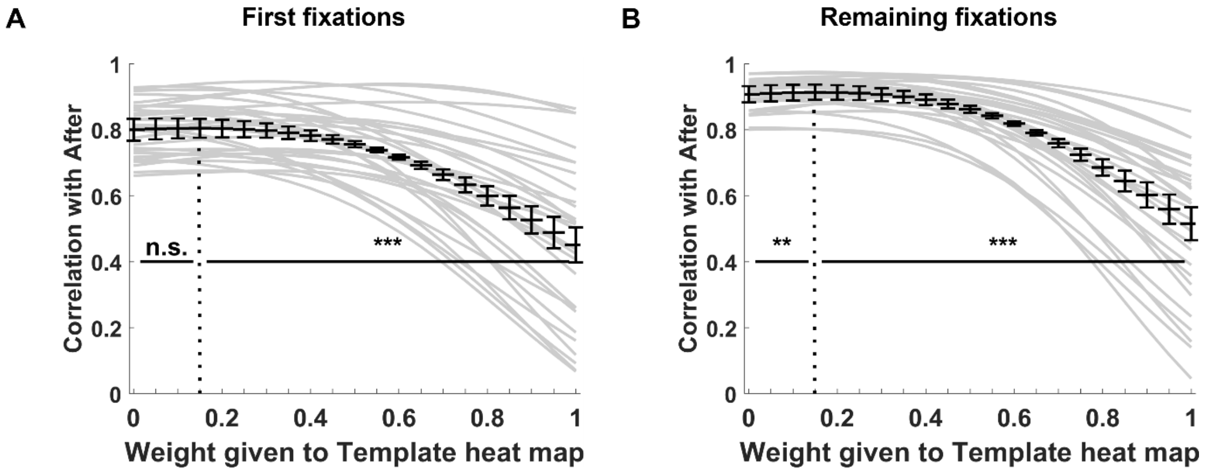


*Fig. S2. Linear combination analysis for first fixations (panel A) and the remaining fixations (panel B) for Experiment 3. Here, the heatmaps from dummy templates, instead of correct templates, were included in the combinations.*

According to our hypothesis about the role of object representations in gaze control, in Experiment 1, in comparison to Experiment 3, the information from templates should play a larger role in driving gaze behaviour in the After condition because only in this experiment the correct templates were presented. Although the differences in the optimal weights between experiments already suggested that this is the case, we also tested this hypothesis directly. For each image, we compared the optimal weights assigned to template heatmaps between Experiments 1 and 3; we expected them to be higher in the former case. The average of these values amounted to 0.39 (SD = 0.28) for first fixations and to 0.64 (SD = 0.18) for the remaining ones in Experiment 1, and, respectively, to 0.23 (SD = 0.24) and 0.32 (SD = 0.33) in Experiment 3. Both for first and remaining fixations, the difference between the experiments was statistically significant and had the expected direction (first fixations: $t(29) = 2.48$, $p = 0.019$; $M_{diff} = 0.16$, 95% CI = [0.03, 0.29]; remaining fixations: $t(29) = 5.20$, $p < 0.001$; $M_{diff} = 0.33$, 95% CI = [0.20, 0.46]). Together, these results corroborate that in the After condition, gaze behaviour is influenced by prior object-knowledge.

**Data exclusion**

Each observer viewed each image in each condition for 3 seconds. Some of these trials were discarded from our analyses because of the low amount of recorded data (Table S1). Specifically, we first excluded all trials during which no fixations were registered (e.g., because the observer did not move their gaze from the location of the fixation point). For all remaining trials, we calculated the percentage of the eye-tracker data-samples, in which the eye-position was not recorded (e.g., due to blinking). After visually inspecting histograms of the obtained values, we excluded from further analyses all viewing sessions for which more than 30% of the position data was missing. Meaningfulness ratings for images that were excluded in the Before and/or the After condition were not analysed. The number and percentage of excluded trials is summarized in Table S1.

Table S1. Excluded viewing-session per experiment

| Experiment Number | Number of excluded trials with no fixations | Number of excluded trials with too few data points or no fixations | Number of trials | Percentage of excluded trials |
|---|---|---|---|---|

| | | | | |
|---|---|---|---|---|
| 1 | 12 | 46 | 3240 | 1.79% |
| 2 | 7 | 60 | 1620 | 4.14% |
| 3 | 6 | 13 | 1800 | 1.33% |

**Normalized entropy calculation**

Entropy calculated for a heatmap provides a measure of its spread. This measure is not dependent on the map's shape. However, its values are dependent on the number of fixations used to create the heatmap (Gameiro, Kaspar, König, Nordholt, & König, 2017; Wilming, Betz, Kietzmann, & König, 2011). Given that the heatmaps from our experiments differed with respect to the total number of fixations underlying them, we estimated entropy values by means of a bootstrapping procedure which accounts for these differences (Gameiro et al., 2017). Specifically, for a given image, we first randomly selected 50 fixations from the pool of all fixations, converted them into a heatmap, and calculated its entropy using a standard Matlab function (*entropy*). This procedure was then repeated 50 times and the entropy values obtained in all the iterations were averaged.

The absolute value of a heatmap's entropy also depends on the specific binning of a heatmap, i.e., the range of possible pixel values (Onat, Açik, Schumann, & König, 2014). Because we were interested only in the changes of entropy between conditions, rather than in the absolute values, we normalized the values from the bootstrapping procedure to the range from zero to one – hence the term normalized entropy. The normalization was performed by dividing the entropy values by the maximal entropy-value possible to obtain for a heatmap, given the size of our images and the binning. This theoretical maximal value was calculated as the entropy of a heatmap being a uniform random distribution (without any smoothing).
